# Spatially correlated fluctuations govern relative chromatin motion

**DOI:** 10.64898/2026.03.10.710930

**Authors:** Janni Harju, Mattia Ubertini, Deepthi Kailash, Po-Ta Chen, Pierre Ronceray, Luca Giorgetti, Thomas Gregor, David B. Brückner

## Abstract

Essential nuclear processes require pairs of chromosomal loci to find each other in three-dimensional space. Polymer models of chromosome dynamics typically assume that the stochastic forces driving such locus motion are spatially uncorrelated, implying that relative diffusion follows directly from single-locus dynamics. Here we show that this assumption fails in living cells. Using live-cell imaging in fly embryos and mouse embryonic stem cells, we find that pairwise locus distances diffuse markedly slower than predicted for independent fluctuations. Combining stochastic trajectory analysis with polymer simulations, we demonstrate that this slowdown arises from non-equilibrium spatially correlated fluctuations (SCFs) in the nucleoplasm, which cause nearby loci to move coherently. We establish three experimentally testable signatures of SCFs: fluctuation amplitudes plateau at large distances, are independent of genomic separation, and show an anomalous temporal scaling. All three predictions are confirmed experimentally, including for loci on separate chromosomes. ATP depletion and disruption of cohesin-mediated loop extrusion reveal that both active processes and crosslinking contribute to correlation magnitudes. Because SCFs slow relative motion preferentially at short distances, they reduce encounter frequencies while prolonging encounter durations, generating a trade-off with direct implications for gene regulation. Our results identify spatially correlated fluctuations as a fundamental determinant of relative motion in confined active polymers.

---

From gene regulation to DNA repair, many processes essential to cellular life share a common biophysical problem: pairs of DNA elements must find each other in three-dimensional space. Enhancers must physically contact their target promoters to activate transcription, often across tens to hundreds of kilobases [1–5]. The ends of a double-strand break must find each other and remain in proximity long enough for repair [6]. How far two DNA loci must travel to meet depends on chromosome organization: chromosome conformation capture methods have revealed hierarchical structures, including compartments [7, 8] and topologically associated domains (TADs) [9–11]. Yet, live-cell imaging reveals that this organization is dynamic: TADs assemble and disassemble on time-scales of minutes [12–14], micron-scale chromatin regions exhibit coherent flows [15, 16], and tagged DNA loci explore large spatial regions through subdiffusion [1, 17–19]. To understand how DNA loci find each other, we must therefore characterize the dynamics of their relative motion.

The encounter frequencies of two loci are governed by the subdiffusive dynamics of their distance. A foundational assumption of polymer dynamics is that the stochastic forces driving monomer motion are uncorrelated in space and time. Under this assumption, at time-scales shorter than those required for stress propagation along the connecting polymer segment, relative diffusion follows directly from the sum of independent single-locus dynamics. This spatial independence underlies most polymer models of chromatin, from the classical Rouse model to more recent theories and simulations incorporating loop extrusion and other processes [20–25].

Here, we show that this assumption fails in living systems. We reveal that spatially correlated fluctuations (SCFs) substantially alter relative diffusion, causing pair-wise distance dynamics to deviate strongly from predictions based on independent noise. Combining live-cell imaging and stochastic trajectory analysis in fly embryos and mouse embryonic stem cells, we demonstrate that relative diffusivities can decrease by more than 50% when loci are in close spatial proximity. Theory and simulations indicate that this dependence arises from SCFs mediated by the nucleoplasm: nearby loci experience similar local forces and thus move coherently. This invalidates the spatial independence of stochastic forces assumed in most polymer models. We test the key predictions of this framework experimentally, showing that SCFs also couple loci on separate chromosomes and that their magnitude decreases upon depletion of ATP or loop-extrusion factors. Finally, our model predicts that SCFs reduce the frequency of pairwise locus encounters while prolonging their duration, generating a trade-off with functional implications for gene regulation.

## I. LOCUS PAIR DISTANCES DIFFUSE ANOMALOUSLY SLOWLY

To investigate the stochastic dynamics of pairwise chromosomal locus motion, we analyze live-cell fluorescence microscopy data from fruit fly embryos [18] (SI Appendix 2.1). Two DNA loci – an enhancer and a genetically inserted promoter paired with a *homie* insulator element – were tracked at variable genomic separations *s* (Fig. 1c). The diffusion of a single locus with position **r**(*t*) at time *t* is quantified by the mean-squared displacement (MSD), which scales as MSD(*t*) = Γ_1_*t*^*β*^, where Γ_1_ is the single-locus subdiffusive coefficient, and *β*≈ 0.5 is the subdiffusive exponent [18]. For two loci *i* and *j*, their displacement vector **R** = **r**_*i*_ − **r**_*j*_ shows similar subdiffusive behavior at short time-scales: MSD_2_(*t*) ≈ 2Γ_2_*t*^*β*^, where Γ_2_ is the two-locus subdiffusive constant, before plateauing at the mean-squared distance MSD_2_ ≈ 2 ⟨*R*^2^⟩ (Fig. 1a). Strikingly, the intercepts of the two-locus MSD curves, which determine 2Γ_2_, are systematically below the value 2Γ_1_ expected if the loci were moving independently. This deviation depends strongly on *s*: locus pairs at shorter genomic separations diffuse markedly slower than those at larger separations (Fig. 1b, SI Appendix 5.1, 5.2). Notably, this reduction is also observed in separate experiments without the *homie* insulator element, indicating that the effect is not sensitive to the local genomic context of the DNA loci (SI Appendix Fig. S12). The assumption Γ_2_ = Γ_1_ is therefore violated across all probed genomic separations and contexts.

**FIG. 1.**
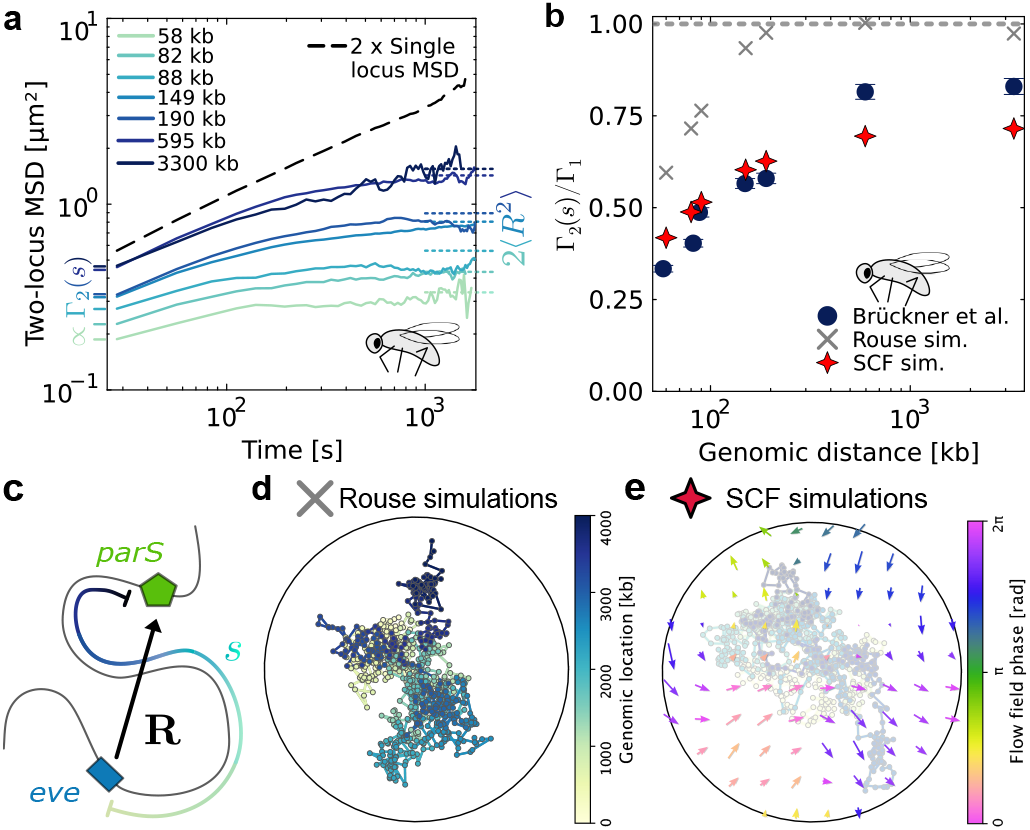
Locus pair diffusion. **(a)** Experimental two-locus MSD curves, 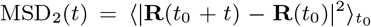 for a range of *s* values. Dashed black line indicates prediction before plateau levels if locus pairs moved independently, MSD_2_(*t*) = 2MSD(*t*). Y-intercepts are proportional to Γ_2_(*s*). Dashed colored lines on the right indicate the plateau levels ⟨*R*^2^⟩. **(b)** The two-locus MSD coefficient Γ_2_ relative to the single locus MSD Γ_1_, for the experimental data and Rouse simulations with or without SCFs. Errors bars calculated from s.e.m. of ⟨Δ*R*^2^⟩ and ⟨Δ*r*^2^⟩. Error bars for simulated data negligible. **(c)** We track the motion of two fluorescently labeled loci inside living *D. melanogaster* embryo cells [18], separated by a varied genomic separation *s*. **R** is the distance between the loci. **(d)** Illustration of 2D Rouse simulations without spatial noise correlations. Circle depicts confinement. **(e)**Illustration of 2D SCF simulations. Arrows depict correlated flow field. Unless otherwise stated, we show 2D simulation results. Results are similar in three dimensions (SI Appendix, Fig. S6).

What could cause this reduction? For an imaging interval Δ*t*, the two-locus diffusivity is determined by the covariance of the distance fluctuations, 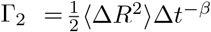, where Δ*R* = |**R**(*t* + Δ*t*) − **R**(*t*) | [25, 26]. Thus, Γ_2_ can be decomposed as (SI Appendix 4.1):

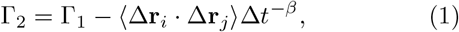

where Δ**r** is the locus displacement between frames. The first term gives the diffusivity assuming independent locus motion; the second captures correlated motion, which suppresses relative fluctuations. The observed reduction of Γ_2_ therefore implies correlated motion between loci. This raises a central question: what is the origin of these correlations?

## II. SPATIALLY CORRELATED FLUCTUATION MODEL RECAPITULATES TWO-LOCUS DIFFUSION BEHAVIOR

In principle, such correlations could arise from two distinct mechanisms. First, they could be mediated through the polymer segment connecting the loci and depend solely on the genomic separation *s*. In any polymer model with reciprocal interactions and uncorrelated noise acting on each bead, such as the Rouse model, center-of-mass motion of the connecting segment introduces correlations that scale as *s*^−1^ [25]. However, the experimental data do not follow this scaling (SI Appendix, Fig. S12a), and Rouse simulations matched to experimental length and time scales exhibit a substantially weaker *s*-dependence of Γ_2_ than observed (Fig. 1b, SI Appendix 3, Fig. S4). Purely polymeric correlations are therefore insufficient to explain the data.

Second, correlations could be mediated through three-dimensional space, for example, via hydrodynamic interactions, active flows, or cross-linking. Such spatially correlated fluctuations (SCFs) are generic in soft matter, from colloidal suspensions [27, 28] to active gels [29], and would generate correlations that depend on the spatial distance *R* rather than the genomic separation *s*.

Motivated by experimental observations of spatially coherent chromatin motion [15, 16] and previous modeling work [30, 31], we introduce a minimal SCF model, consisting of a Rouse chain immersed in a random, divergence-free flow field, consistent with incompressible flows. This flow field is characterized by an average magnitude *u*_flow_, a correlation length *λ*_flow_, and a correlation time-scale *T*_flow_ (SI Appendix 3, Video S1). Our simulations predict Γ_2_(*s*) curves consistent with experiments (Fig. 1b). Intuitively, although SCFs depend on the spatial distance *R* rather than *s*, locus pairs at shorter genomic separations are more frequently spatially proximal and hence experience stronger correlations. Non-equilibrium, flow-mediated SCFs hence recapitulate the experimental data by introducing a distance-dependence into the two-locus diffusivity.

## III. STOCHASTIC TRAJECTORY ANALYSIS REVEALS SPATIAL CORRELATIONS

Our SCF model makes a key prediction: if correlations are spatial, the magnitude of pairwise distance fluctuations should depend on the instantaneous three-dimensional distance *R*, rather than the genomic separation *s*. To test this directly using experimental trajectories, we define the fluctuation-distance profile

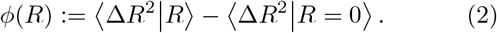

This quantifies how distance fluctuations increase when the loci are farther apart relative to the fluctuation magnitude at *R* = 0.

Even without SCFs, the Rouse model predicts a non-trivial *ϕ*(*R*): conditioning on a large initial distance *R* selects extended polymer configurations whose relaxation increases fluctuations. Intuitively, the excess mode amplitudes scale with *R*, and over time-scales Δ*t* shorter than the segment’s relaxation time *τ*_*s*_, these modes relax by a fraction ∼ Δ*t/τ*_*s*_. The polymeric contribution to *ϕ* can then be shown to scale as *ϕ*_Rouse_ ∼ *R*^2^*s*^−2^Δ*t* (derivation in SI Appendix 4.2).

By contrast, if SCFs with a decaying spatial correlation function such as *C*(*R*) = *C*_0_*e*^−*R/λ*^ dominate, we find (SI Appendix 4.2):

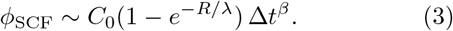

The two models thus yield three distinguishing and experimentally testable signatures: (i) while the Rouse model predicts convex (∼ *R*^2^) scaling, SCFs predict a concave plateau, (ii) the Rouse scaling depends on the genomic separation through *s*^−2^, whereas SCFs predict *s*-independence, and (iii) the Rouse model predicts *ϕ* ∼ Δ*t*, whereas SCFs predict *ϕ* ∼ Δ*t*^*β*^. These qualitative signatures do not depend on the precise form of the correlation function *C*(*R*), as long as it decays in a convex manner with distance.

We first validate these predictions in simulations, focusing on genomic separations *s* > 100 kb where SCFs dominate over polymeric effects at the experimental imaging interval Δ*t* = 28 s (SI Appendix 4.3). In the absence of correlated flows, *ϕ* scales as *R*^2^ and collapses when plotted as *ϕ*Δ*t*^−1^ against *Rs*^−1^, as expected for purely polymeric correlations (Fig. 2a,e, SI Appendix 4.4, Fig. S8, S10). With correlated flows, by contrast, *ϕ* plateaus at large *R*, is independent of *s*, and scales with Δ*t*^*β*^ at short distances (Fig. 2b,f, SI Appendix Fig. S10). For *s <* 100 kb, SCF simulations show the predicted crossover to Rouse-like scaling at large distances (SI Appendix Fig. S7).

**FIG. 2.**
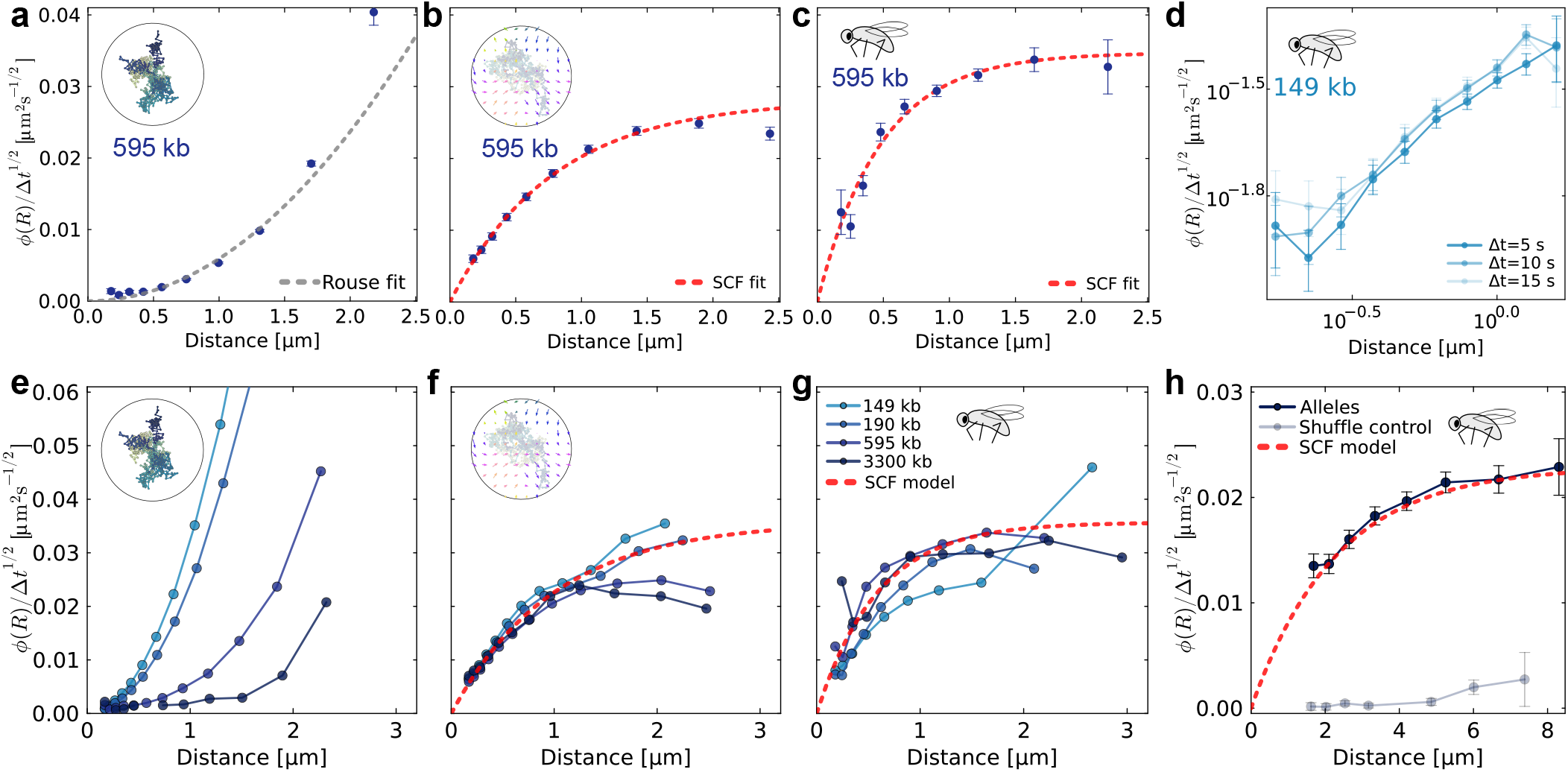
Fluctuation-distance profiles reveal spatial correlations. **(a)** Spatial dependence of fluctuation-distance profile for *s* = 595 kb in Rouse simulations. Gray line shows best fit for quadratic scaling. **(b-c)** Similar to (a), but for SCF simulations and experimental data from [18]. Red line shows best fit for 1 − *e*^−*R/λ*^ scaling. **(d)** Collapse of *ϕ* for experimental data sub-sampled at different Δ*t*. **(e)** Plots of *ϕ*/Δ*t*^1/2^ at fixed Δ*t* ≈ 28 s for simulations of a Rouse model, for varying *s*. The data can be collapsed for *Rs*^−1^ (SI Appendix Fig. S8). **(f-g)** Similar to (e), but for SCF simulations and experimental data from [18]. Red line indicates expected SCF scaling with 1 − *e*^−*R/λ*^. Curves for lower *s* start to curve upwards at larger *R*, consistent with a transition into the Rouse scaling regime (SI Appendix 4.3). All individual curves in panels e-g are shown with error bars in SI Appendix Fig. S7. **(h)** *ϕ*/Δ*t*^1/2^ curves for tracked inter-chromosomal loci, obtained by tracking two alleles of the gene *Krüppel* (SI Appendix 2.2). Grey line corresponds to data shuffled between nuclei, where no locus motion correlations are expected. In all panels, error bars indicate s.e.m. for each bin.

Analysis of our fly embryo data confirms all three predicted SCF signatures: fluctuation-distance profiles *ϕ*(*R*) show no systematic dependence on *s* (Fig. 2g), plateau at large *R* for *s* > 100 kb (Fig. 2c, g), crossover to Rouse-like behavior for *s <* 100 kb (SI Appendix S7), and collapse with Δ*t*^*β*^ across subsampled time intervals (Fig. 2d). These findings are robust to variations in bin size (SI Appendix Fig. S11) and to the presence or absence of *homie* insulator elements at the tracked loci (SI Appendix Fig. S12c). By fitting Eq. 3 to the experimental fluctuation-distance profiles, we estimate a correlation length of *λ* ≈ 0.6 μm (SI Appendix 5.3, Table S4). Taken together, our results show strong evidence that SCFs define two-locus motion in living fruit fly embryos.

## IV. SCFs COUPLE INTER-CHROMOSOMAL LOCUS MOTION

Our SCF model makes a key prediction: since correlations are mediated through the nucleoplasm rather than the polymer backbone, they should also couple the motion of loci on separate chromosomes. To test this, we experimentally tracked the distance between the two allelic copies of the same locus, located on separate chromosomes, in fly embryos (SI Appendix 2.2, Fig. S1, Videos S2, S3). The fluctuation-distance profiles follow the same concave form as for loci on the same chromosome (Fig. 2h). As a control, shuffling cell labels to pair alleles from different nuclei yields flat fluctuation-distance profiles, as expected in the absence of correlated motion (Fig. 2h, SI Appendix 2.2.7). SCFs thus act across chromosomes, consistent with correlations mediated through the nucleoplasm.

## V. SPATIAL CORRELATIONS ARE CONSERVED IN MAMMALIAN CELLS

To test whether SCFs affect locus motion in other systems beyond fly embryos, we analyze two-locus tracking data from mESCs [13, 32] (SI Appendix 2.1). We find that across mESC data from three genomic separations, generated by two independent laboratories at different temporal resolutions, fluctuation-distance profiles match all three predictions of the SCF model: they are concave, independent of *s*, and collapse for *ϕ* ∼ Δ*t*^*β*^ with *β* ≈ 0.5 (Fig. 3a,b, SI Appendix Fig. S10). Furthermore, across species, genomic separations, and genomic perturbations, Γ_2_ scales with ⟨*R*⟩*/D*, the mean interlocus distance normalized by mean nucleus diameter *D* (Fig. 3c, Pearson correlation 0.98), suggesting that spatial distances are the main determinant of two-locus diffusivities across different organisms.

**FIG. 3.**
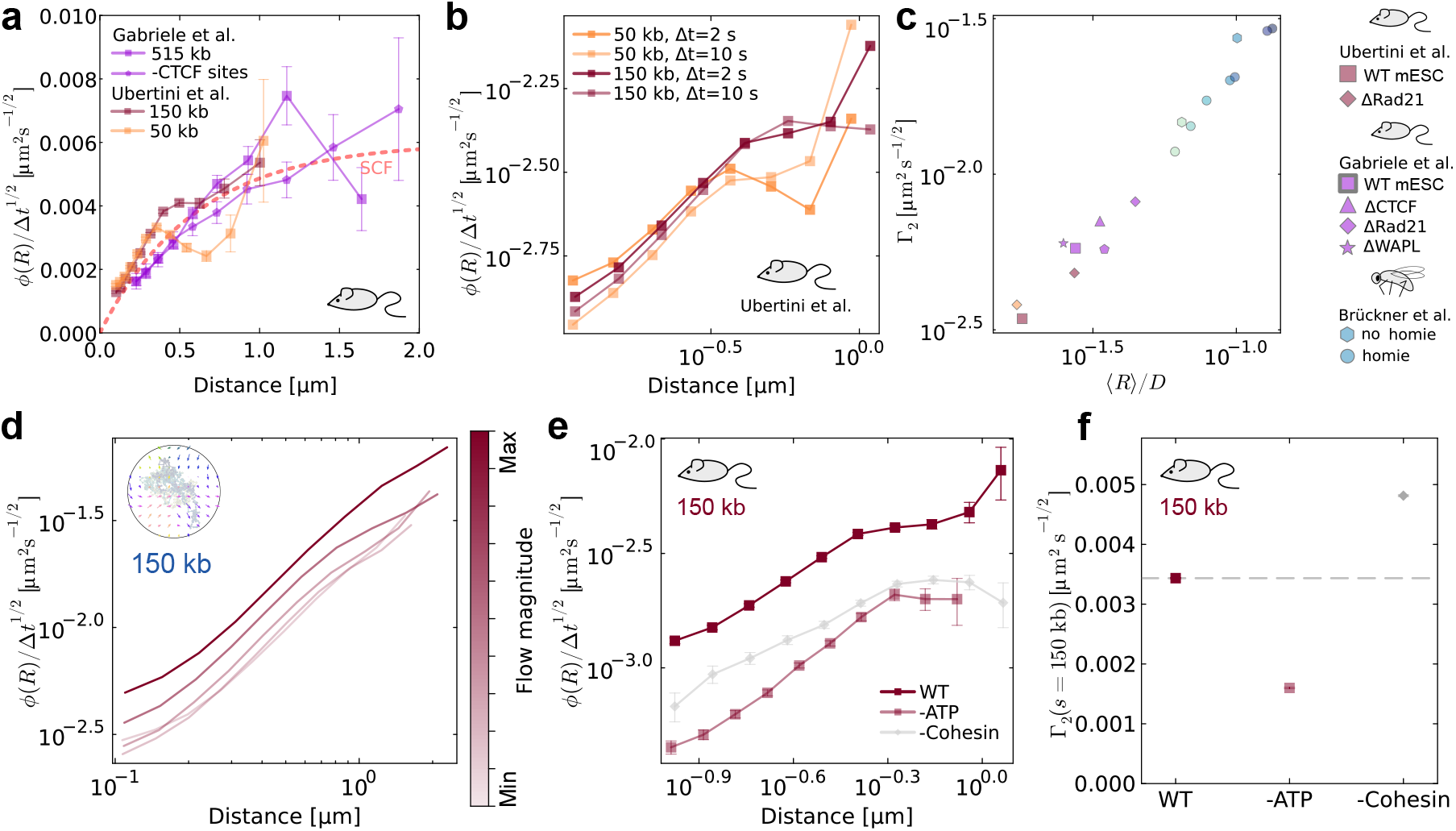
SCFs in mESC data decrease with nuclear activity. **(a)** *ϕ* for mESC data with three genomic separations from [13, 32]. -CTCF sites curve shows that removal of CTCF binding sites at labeled loci does not affect SCFs. Individual fluctuation-distance profiles with error bars are shown in SI Appendix Fig. S13, S14. **(b)** Collapse of *ϕ* for mESC data from [32]. The fluctuation-distance profiles are independent of the genomic separation, and scale with Δ*t*^1/2^, as predicted by the SCF model. **(c)** Scatter plot of Γ_2_ = ⟨Δ*R*^2^⟩ Δ*t*^−1/2^ vs. mean locus distance over cell diameter. Pearson correlation coefficient is 0.98. **(d)** Simulation prediction for effect of decreasing active flow magnitude on *ϕ*. **(e)** Experimental data for effects of ATP depletion (SI Appendix 2.3) and cohesin depletion [32] on fluctuation-distance profiles in mESCs. **(f)** Experimental data for effects of ATP depletion (SI Appendix 2.3) and cohesin depletion [32] on Γ_2_(*s*) in mESCs.

## VI. SCFs ARE ACTIVELY DRIVEN

What mechanisms underlie SCFs? We consider two qualitatively different sources. Active flows, driven by hydrodynamic interactions that cause alignment of motor proteins within the nucleoplasm [33], would both increase SCF magnitudes and locus diffusivities. Chromatin crosslinking or coupling to a surrounding viscoelastic medium, by contrast, would increase SCFs but decrease diffusivities by constraining motion. These opposing predictions for Γ_2_ provide a way to distinguish between the two types of mechanisms experimentally. Furthermore, crosslinking and viscoelastic couplings could give rise to passive, rather than active, spatial correlations.

Our model recapitulates the experimental data using active flows, and an alternative implementation of SCFs as a spatially correlated but temporally uncorrelated equilibrium noise field [34] (SI Appendix 3, Video S4) is quantitatively inconsistent with experimental fluctuation-distance profiles (SI Appendix 4.5). To experimentally test whether activity gives rise to flow-like correlations in living systems, we deplete ATP in mESCs (SI Appendix 2.3, Fig. S3, Videos S5, S6). Our simulations predict that reducing the magnitude of active flows decreases both *ϕ*-magnitudes and Γ_2_ (Fig. 3d, SI Appendix Fig. S5b). Consistently, ATP depletion in experiments also decreases *ϕ*-magnitudes and Γ_2_ (Fig. 3e,f), suggesting reduced flow-mediated correlations. This demonstrates that active, ATP-dependent processes contribute to SCFs.

## VII. LOOP EXTRUSION CONTRIBUTES TO SCFs VIA CROSSLINKING

A particularly relevant active process for chromosome dynamics is loop extrusion by cohesin, which actively reels in chromatin loops during interphase [10, 35]. Perturbing loop extrusion could affect pairwise dynamics in at least three ways: by loosening chromosome organization and thus increasing the mean distance between loci [13, 32]; by removing crosslinks that couple the motion of nearby loci; or by reducing motor-driven flows within the nucleoplasm.

Consistent with previous reports [12, 13, 32], we find that cohesin depletion increases both mean interlocus distances ⟨*R*⟩ and Γ_2_ (Fig. 3e, f). These increases follow the overall scaling of Γ_2_ with ⟨*R*⟩*/D*, suggesting that much of the change in two-locus diffusivities reflects the change in mean distance. To disentangle the effects of the change in mean distance from possible changes in correlation magnitudes, we calculate *ϕ*(*R*) under cohesin depletion. The decrease in the magnitude of *ϕ*(*R*) (Fig. 3e) indicates weaker correlations at fixed spatial distances. This combination of decreased correlation magnitudes and increased diffusivities points to a crosslink-like mechanism: cohesin removal both reduces correlation magnitudes and frees individual loci to move faster.

## DISCUSSION

In this work, we showed that spatially correlated fluctuations within the nucleoplasm are a major determinant of the relative motion of chromosomal locus pairs. Using theory and simulations, we established three signatures of the fluctuation-distance profile *ϕ*(*R*) that distinguish SCFs from purely polymeric correlations: concave *R*-dependence, independence from genomic separation, and a Δ*t*^*β*^ temporal scaling. Experimental data from fly embryos and mESCs match all three signatures. Our experiments demonstrate that SCFs couple loci on separate chromosomes, and that their magnitude depends on both active energy-dependent processes and cohesin-mediated crosslinking. Together, these results identify spatially correlated fluctuations as a previously unrecognized physical driver of chromosome dynamics.

Biologically functional interactions between DNA locus pairs depend on two time-scales: the first-passage time of the loci, which sets the interaction frequency [36, 37], and the encounter duration, which determines whether an interaction can initiate transcription [32]. Our model predicts that SCFs create a trade-off between these two time-scales (Fig. 4). Because correlated motion slows relative diffusion at short distances, encounters between nearby loci become less frequent but longer-lived (Fig. 4a,b).

**FIG. 4.**
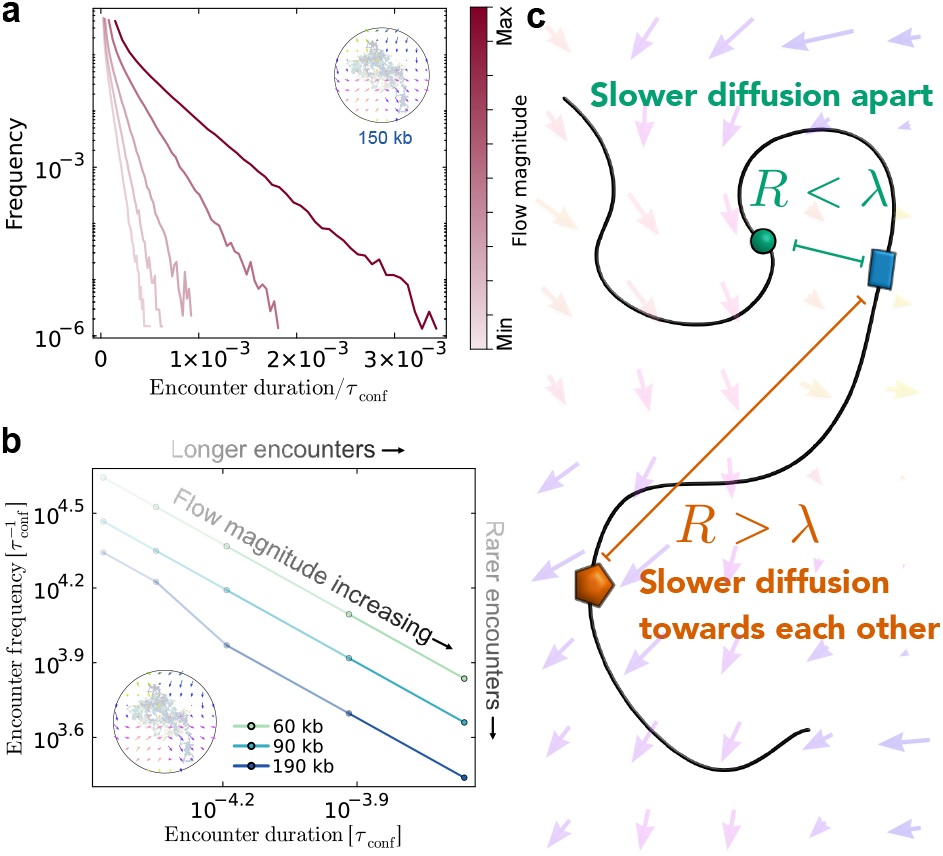
Spatial correlations increase duration and decrease frequency of encounters. **(a)** Frequency of encounter durations (periods of time for which *R < r*_enc_) for simulations with varying flow magnitudes. **(b)** Encounter frequency vs. duration from SCF simulations. Line opacity reflects mean flow field magnitude *u*_flow_, values same as in panel (a). Both axes are plotted in units of *τ*_conf_ = (*D*^2^/Γ_1_)^2^, the time it takes for a single locus to diffuse across the entire confinement of diameter *D*. **(c)** Schematic of effects of SCFs on encounter time-scales. Arrows illustrate a correlated flow field. SCFs increase encounter durations, but also decrease encounter frequencies.

This trade-off has direct implications for enhancer-promoter communication: fewer encounters may reduce transcription initiation rates, whereas longer encounters provide more time for the assembly of transcriptional machinery once contact is established (Fig. 4c). While loop extrusion prolongs encounters, and may thereby increase the probability of transcription initiation [32], SCFs provide an additional, independent mechanism for prolonging encounters that persists even after loop-extrusion factor degradation. Given that chromatin dynamics are strongly subdiffusive [19], SCFs further slow an already constrained search process, amplifying their impact at short distances where regulatory interactions occur. Beyond transcription, SCFs are expected to influence processes in which relative polymer motion is rate-limiting, including DNA repair [6, 38], meiotic chromosome pairing [39, 40] and RNA transport and splicing [41–43].

The physical origin of SCFs likely involves multiple mechanisms, which will need to be resolved using further simulations and experiments. We find evidence that loop-extrusion factors contribute to SCFs via a crosslink-like mechanism. Additional contributions may arise from passive effects such as hydrodynamic correlations [27, 28] or viscoelastic couplings, for example, at the nuclear periphery [44, 45]. Correlations may further be modulated by nuclear condensates and compartments [46], which locally alter the effective viscosity of the medium [47, 48]. Finally, chromatin constitutes an active polymer melt in which molecular motors and other energy-consuming processes generate forces that propagate through the surrounding nucleoplasm [33, 49]. Two-fluid models treating chromatin and solvent as coupled interpenetrating fluids show how distributed activity can generate coherent flows [33, 50], while other theoretical approaches consider active forces correlated along the genomic coordinate [51– 53]. Polymer simulations in imposed flow fields have explored effects on chain conformations [30, 31], but have not addressed pairwise locus dynamics. Here, we demonstrate that active flows have dramatic consequences for the pairwise dynamics of chromosomal loci.

Long-range velocity correlations are a generic feature of active polymers. We show that such correlations fundamentally alter the relative motion between polymer segments, challenging a foundational assumption underlying most polymer models for chromatin. More broadly, this work provides a framework for detecting, quantifying, and interpreting spatial correlations in active polymer systems across biological contexts.

## ACKNOWLEDGMENTS

We thank Miloš Nikolić, Alex Chen Yi Zhang, and Chase Broedersz for discussions and feedback. We would like to thank Anupama Hemalatha for kindly sharing rotenone and 2-deoxy-D-glucose. This work was supported by the Princeton Presidential Postdoctoral Fellowship and the Center for the Physics of Biological Function (JH); the EMBO Postdoctoral Fellowship (ALTF 1305-2024, MU); the Novartis foundation and the Swiss National science Foundation (310030 192642, TMCG-3 213782 and 320030-236070, LG); the French National Research Agency (ANR-20-CE12-0028 ‘ChroDynE’ and ANR-23-CE13-0021 ‘GastruCyp’, TG), the European Research Council (ERC-2023-SyG, ‘Dynatrans’, 101118866, TG), the U.S. National Science Foundation, through the Center for the Physics of Biological Function (PHY-1734030, TG), and by the National Institutes of Health (R01GM097275, U01DA047730, and U01DK127429, TG); and support from the Biozentrum (DBB).

## AUTHOR CONTRIBUTIONS

D.B.B. conceived the project. J.H., P.R., D.B.B. proposed the model. J.H. developed theory and analyzed data. J.H developed and performed simulations with input from P.R. and D.B.B.. M.U. performed mESC experiments and the corresponding image analysis. P.-T.C. performed fly experiments. D.K. performed image analysis for fly experiments. L.G. and T.G. supervised experimental and data analysis work. J.H. and D.B.B. wrote the paper with input from all authors.

## DATA AVAILABILITY

Previously published experimental data used in this study can be found at [54–56]. Allele tracking data for fruit fly embryos can be found at 10.5281/zenodo.18930334.ATP depletion data for mESC cells can be found at 10.5281/zenodo.18912187. Simulation data can be found at 10.5281/zenodo.18938320.

## CODE AVAILABILITY

The simulation and analysis code used in this study can be found at github.com/harjuj/Chromatin_SCFs.

## METHODS

### Fly embryo experiments

We analyzed previously published two-locus tracking data from *Drosophila melanogaster* embryos [18], in which an endogenous *eve* locus (MS2-labeled) and a reporter promoter (*parS* - labeled) were imaged at 28 s intervals across a range of genomic separations *s*.

To test whether spatially correlated fluctuations (SCFs) couple loci on separate chromosomes, we performed new two-photon imaging experiments tracking two allelic copies of the *Kr* locus, labeled with MS2 and MS2-PP7 stem loops, in fly embryos at nuclear cycle 14 (SI Appendix, Fig. S1). Stacks of 12 *z*-planes (1 μm spacing) were acquired every ∼10 s using 920 nm and 1045 nm excitation channels. Nuclei were segmented with a custom-trained Cellpose model [57] and tracked via nearest-neighbor assignment. Gene loci were detected in each z-slice using adaptive non-local means filtering [58], stitched along z, and localized by intensity-weighted centroid. Locus pairs were matched across channels and tracked over time. Based on the experimentally derived point spread function, we estimated a minimum resolvable distance of *σ*_*R*_ = 1.43 μm and restricted analysis to *R > σ*_*R*_ (SI Appendix, Fig. S2). As a control, we shuffled allele tracks across nuclei to construct pairs expected to show no correlated motion. For more details, see SI Appendix Section 2.2.

### Mouse embryonic stem cell (mESC) experiments

We analyzed previously published mESC data sets in which pairs of loci were labeled with TetO/LacO or TetO/Anchor3 arrays at genomic separations of 50– 515 kb and imaged at 2 s [32] or 20 s [13] intervals. Perturbations included auxin-induced degradation of RAD21, CTCF, and WAPL.

To test whether SCFs depend on active processes, we acquired new data in a previously established mESC line carrying TetO and LacO arrays separated by 150 kb [12]. ATP was depleted by treating cells with 1 μM rotenone and 6 mM 2-deoxy-D-glucose for 1 h prior to imaging [59]. Cells were imaged in oblique illumination mode on a Nikon Eclipse Ti2 widefield microscope at 0.5 Hz (2 s frame interval) for 5 min, acquiring 9 z-planes spaced 0.3 μm apart. Spot detection and tracking followed the pipeline described in [32]. Tracks with anomalously large frame-to-frame displacements (*>* 200 nm^2^) were manually inspected and filtered to remove artifacts from replicated alleles (SI Appendix, Fig. S3). For more details, see SI Appendix Section 2.3.

### Data analysis

Single-locus subdiffusive coefficients were calculated as 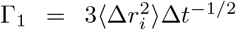from the *i*-components of frame-to-frame displacements, and two-locus coefficients as 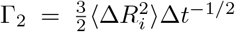, from the *i*-components of frame-to-frame changes in the inter-locus vector **R**. Only *x* or *y* displacements were used for experimental data, due to the larger localization error in the *z*-direction.

To quantify the distance-dependence of fluctuations, we defined the fluctuation-distance profile *ϕ*(*R*, Δ*t*) = ⟨Δ*R*^2^ |*R*⟩−⟨Δ*R*^2^ |*R*⟩ = 0 . Conditional averages ⟨Δ*R*^2^| *R*⟩ were computed by logarithmically binning instantaneous distances, discarding data below the experimental resolution. Edge bins with fewer than 30 samples were merged. The binned data were fitted with two models: an SCF prediction ⟨Δ*R*^2^ | *R*⟩ = *A* + 2*C*(1 − *e*^−*R/λ*^), and a Rouse prediction ⟨Δ*R*^2^ | *R*⟩ = *A*^′^ + *BR*^2^, using weighted least-squares fitting. The value ⟨Δ*R*^2^ | 0⟩ was extracted from the better-fitting model evaluated at *R* = 0.

Encounter frequencies and durations were calculated from simulated distance trajectories using Kaplan-Meier survival analysis, with encounters defined as 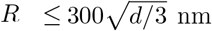, where *d* is the dimensionality of the system. For more details, see SI Appendix Sections 5.3 and 5.4.

### Simulations

We simulated a confined 3D Rouse polymer of *N* = 800 beads (corresponding to 5 kb each) within a harmonic confinement of diameter *D* = 2.8 μm. The spring stiffness *k* and friction constant *ζ* were fixed by matching the experimental mean locus distances and single-locus diffusivities. We considered three models: an uncorrelated Rouse model with independent white noise; (ii) an equilibrium model with spatially correlated white noise; and (iii) a non-equilibrium model where a divergence-free, random flow field advects the polymer. All simulations were implemented in Julia, with GPU acceleration (CUDA.jl) for the equilibrium correlated-noise model. For more details, see SI Appendix Section 3. For all parameter sets, we performed 5-8 independent simulations, each sampled 100 times at 28 s intervals.

The equilibrium simulations with correlated noise included the appropriate noise-induced drift to preserve the equilibrium distribution. The free parameters *f*_cor_ (fraction of correlated noise) and *λ*_cor_ (correlation length) were set by matching Γ_2_(*s*) curves. For all reasonable parameter choices, *ϕ*(*R*) curves did not show SCF scalings at experimentally used time-scales (SI Appendix 4.5).

The non-equilibrium flow field was generated by convolving white noise with a Gaussian filter in the Fourier domain, evolving mode amplitudes as an Ornstein-Uhlenbeck process, and projecting to a divergence-free space. After introducing the flow, *ζ* and *u*_flow_ (mean flow magnitude) were rescaled to maintain the experimental Γ_1_. Many sets of free parameters *u*_flow_, *λ*_flow_ (spatial correlation length), and *T*_flow_ (correlation time) were found to reasonably match both the experimental Γ_2_(*s*) and *ϕ*(*R*) curves. Most flow simulations were performed in 2D for computational efficiency; 3D simulations yielded comparable results (SI Appendix S6).

## Notes

### Competing Interest Statement

The authors have declared no competing interest.

