## Supplementary information for "Spatially correlated fluctuations govern relative chromatin motion"

### Supplementary Information:
#### Spatially correlated fluctuations govern relative chromosomal locus dynamics

Janni Harju, Mattia Ubertini, Deepthi Kailash, Po-Ta Chen, Pierre Ronceray,
Luca Giorgetti, Thomas Gregor, David B. Brückner

**6 Supplementary Tables 40**

**7 Supplementary references 41**

### 1 Supplementary Video Captions

**Video S1.** Animation comparing 2D polymer simulations of a Rouse model (left) with a Rouse model in a spatially and temporally correlated flow field (right). Color of chain indicates genomic coordinate. The correlated flow field is shown with arrows. Color of arrows reflects their orientation. Simulations were conducted in confinement (circle drawn in animations). 3D simulations were also conducted (Fig. S6).

**Video S2.** Maximum intensity projection of merged 920 nm channel and 1045 nm channel during NC14 of the fruit fly embryo. Temporal resolution is 10 s. Scale bar indicates 10 µm.

**Video S3.** Example fruit fly embryo loci imaged in 920 nm channel. Red outline shows the tracked nucleus during NC14. Temporal resolution is 10 s. Scale bar indicates 2 µm. Image is lightly contrasted for visualization purposes.

**Video S4.** Animation comparing 3D polymer simulations of a Rouse model (left) with a Rouse model with equilibrium, spatially correlated white noise (right). Color of chain indicates genomic coordinate. Simulations are conducted in a spherical confinement potential.

**Video S5.** Representative maximum intensity projection of dual-color live cell imaging movie of LacO (green) and TetO (orange) arrays in an untreated mESC. The movie is acquired at 2 s frame interval.

**Video S6.** Representative maximum intensity projection of dual-color live cell imaging movie of LacO (green) and TetO (orange) arrays in a mESC treated for ATP depletion. The movie is acquired at 2 s frame interval.

### 2 Supplementary experimental methods

#### 2.1 Description of previously published data sets

Here, we provide brief summaries of the previously published experimental data sets analyzed in this study. We note that since our analysis requires the calculation of 3D distances separating loci, 2D data sets, such as those recently published in [1], could not be included.

**Brückner *et al.*** In [2], two-locus tracking was performed in *Drosophila melanogaster* embryos. The endogeneous *eve* gene was labeled with MS2 stem loops. A reporter promoter labeled with a *parS* sequence was placed a variable genomic separation $s$ away from *eve*. The imaging interval was 28 s. The reporter cassette also contained a *homie* insulator, which forms stable loops with the endogeneous *homie* locus near *eve*. Control data were obtained without a *homie* insulator near the promoter (Fig. S12c). Importantly, the key predictions of our theory hold for both *homie* and *no-homie* data. For more details on experimental methods for this data set, see ref. [2].

**Gabriele *et al.*** In [3], the motions of two loci flanking the *Fbn2* TAD (515 kb) in mESCs were studied. The sites were labeled using TetO and Anchor3 arrays, which were bound by fluorescently labeled TetR-3x-mScarlet and EGFP-OR3. The imaging interval was 20 s. Additionally, the authors collected data after auxin-induced degradation of CTCF, cohesin factor RAD21, and WAPL. For more details on experimental methods for this data set, see ref. [3].

**Ubertini *et al.*** In [4], two-locus dynamics were studied in mESCs. Briefly, TetO and LacO arrays were inserted at a separation of 50 or 150 kb of each other, within an existing 560-kb TAD. These loci could then be tracked using TetR-tdTomato and a weakly binding, eGFP-labeled LacI variant. The imaging interval was 2 s. Additionally, the work includes data from a cell line where the separation between the loci is reduced from 150 kb to 50 kb. RAD21 depletion was also performed in both cell lines. For more details on experimental methods for this data set, see ref. [4].

#### 2.2 Tracking distances for locus copies on separate chromosomes in fly embryos

##### 2.2.1 Genetic Lines and Crossing

A fly line carrying the fluorescent reporters *(yw; His-RFP; nanos >MCP-eGFP, PCP-mCherry)* was crossed to males from a CRISPR-MS2 transgenic line *(Kr-MS2)*. Female progeny from this cross were subsequently crossed to males from a CRISPR-MS2-PP7 transgenic fly line *(Kr-MS2-PP7)* [5]. Resulting embryos either showed two labeled alleles (one allele labeled in green and one allele labeled in both red and green) or those carrying a single labeled allele (one allele in green) (Fig. S1a,b). Only embryos with two labeled alleles were selected for imaging and subsequent analysis.

##### 2.2.2 Sample Preparation

Sample preparation was adapted from [6, 5]. Embryos were collected on agar plates for 2–2.5 hours. Individual embryos were hand-dechorionated on a piece of double-sided

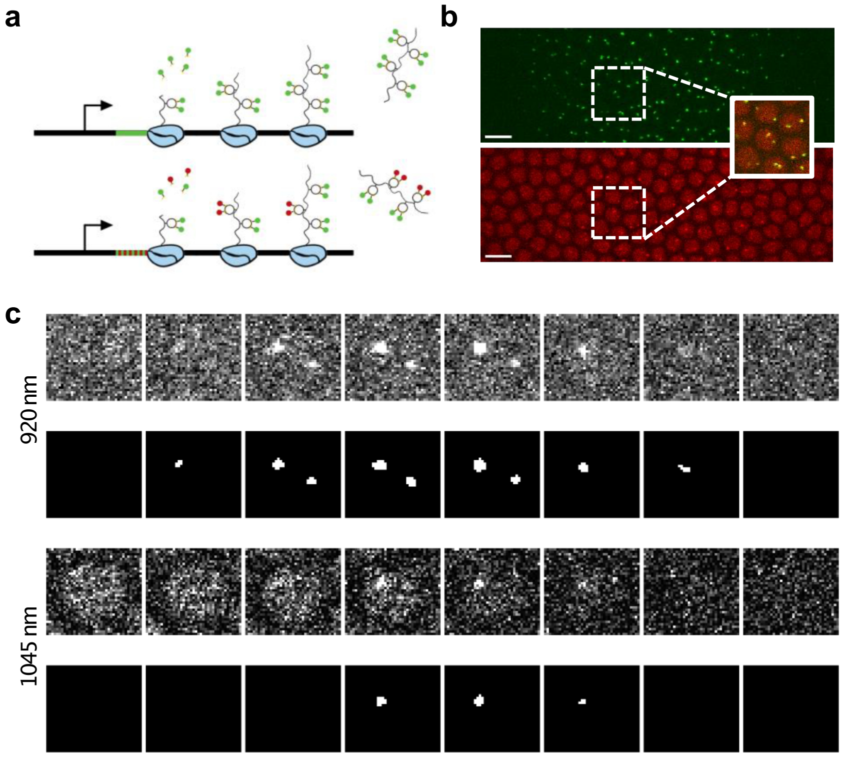

Figure S1: **Experimental setup and analysis of gene loci. (a)** Live imaging of transcription sites using MS2 stem-loops and interlaced MS2-PP7 stem-loops. **(b)** Maximum intensity projections of two imaging channels. The 920 nm excitation channel (top) records two loci per nucleus. The 1045 nm excitation channel records the nuclei and one locus per nucleus. Scale bar is 10 µm. Both images are cropped to the region of active loci. **(c)** Imaging planes and resulting segmentation for a single volume of one nucleus in 920 nm and 1045 nm channels. Planes are spaced 1 µm apart. Localization was done using only the 920 nm channel. Raw images are lightly contrasted for visualization purposes.

tape and then mounted onto a glue-coated membrane. After mounting, embryos were immersed in Halocarbon 27 oil (Sigma) and compressed with a cover glass (Corning #1 1/2, 18 × 18 mm). Excess oil was removed with a tissue, and all embryos were mounted dorsal side up.

###### 2.2.3 Live Imaging

Imaging was performed on a custom-built two-photon microscope as described in [5]. Excitation was achieved using a Chameleon Ultra laser (920 nm, channel 1) and a HighQ-2 laser from Spectra-Physics (1045 nm, channel 2). Excitation and emitted fluorescence were focused and collected through a 40× oil-immersion objective lens (1.3 NA, Nikon Plan Fluor). Laser scanning and image acquisition were controlled using ScanImage (Vidrio Technologies LLC). Fluorescence from both channels was detected simultaneously using separate GaAsP photomultiplier tubes (Hamamatsu H10770P-40). Images were acquired at a pixel size of 220 nm with a frame size of 960 × 540 pixels. Each image stack comprised 12 frames, spaced 1 µm apart along the $z$-axis. Pixel dwell time was 1.4 µs, yielding an overall temporal resolution of approximately 10 s per stack. Embryos were imaged from late NC12 to the end of NC14 (Fig. S1b,c).

###### 2.2.4 Nuclei Segmentation and Tracking

Nuclei were segmented using a custom-trained Cellpose model [7]. For frames after completion of nuclear cycle 13 mitosis, nuclei were tracked across frames using a 3D nearest-neighbor approach. After morphological opening and watershedding to separate nuclei in the mask, nuclear diameter was estimated in 2D from maximum intensity projections by computing an equivalent circular diameter (regionprops). Frames in which an individual nucleus exhibited a measured lateral diameter greater than 10 µm were classified as segmentation errors and excluded from subsequent spot-tracking analyses. Nuclei tracks were verified by visual inspection. All nuclei analysis on the segmented was performed using custom MATLAB scripts.

###### 2.2.5 Spot Segmentation and Tracking

Following manual selection of the time-lapse interval after completion of NC13 mitosis, the original images were partitioned into stacks containing individual nuclei. At each z-slice and time point, gene loci in both fluorescence channels were segmented using an adaptive non-local means filter [8]. The intensity threshold was selected manually per dataset. After an initial detection pass, a weaker threshold was applied to the three z-slices above and below each detected locus to ensure complete inclusion. Detected objects were stitched along the $z$-axis, and a weighted centroid was computed for each locus. This procedure was performed independently for the red and green channels (Fig. S1c).

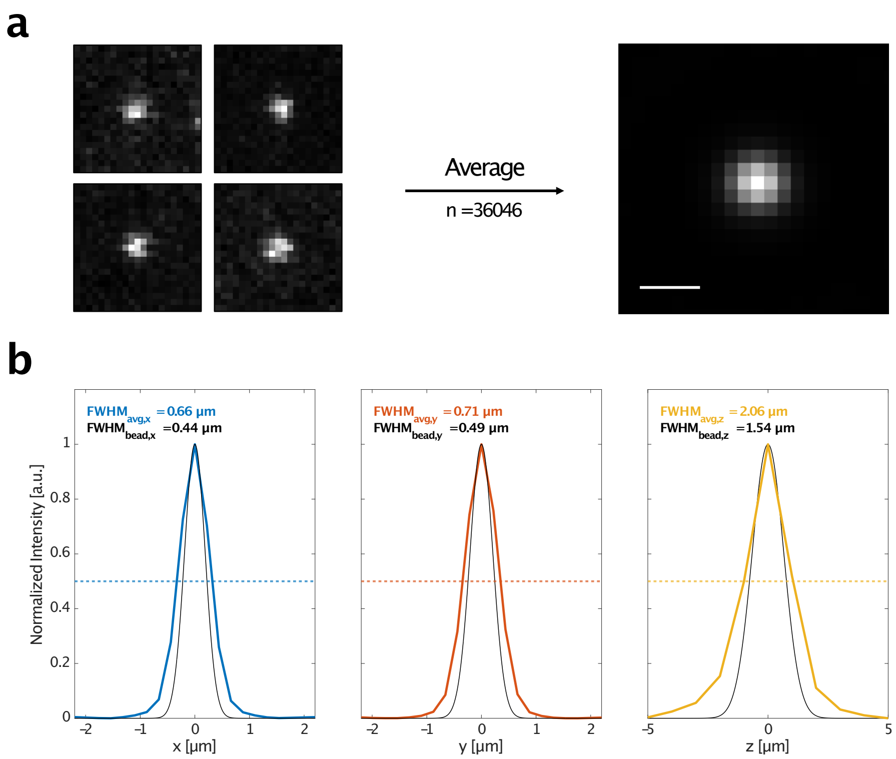

Figure S2: **Distance error estimation using gene loci. (a)** (Left) Maximum intensity projections of four representative gene loci centered within a 21 × 21 × 11 voxel region. A total of 36046 loci were aligned by their centroid and averaged to generate a composite image. (Right) Composite image of loci after background subtraction. Scale bar indicates 1 μm. **(b)** Colors indicate line profiles through the center of the averaged image, with intensity normalized by the maximum. Black lines indicate a gaussian profile with the full-width half maximum from 50 nm beads. Dashed lines are the half maximum in each direction.

To track gene loci over time, loci detected in both channels were first matched across channels at each time point using a distance-based metric. The paired loci were then tracked across frames using a nearest-neighbor approach. Because the signal-to-noise ratio was higher in the 920 nm channel than in the 1045 nm, all centroid and localization measurements were derived from this channel. Frames with only one locus detected or tracks with zero detections in the 1045 nm channel were discarded. Tracks were verified by visual inspection.

##### 2.2.6 Estimating localization error

For each detected spot in each frame across all datasets, a three-dimensional region of interest was extracted by centering a fixed-size subvolume (21 × 21 × 11 voxels) on the spot's intensity-weighted centroid in the XY plane and the Z-plane exhibiting maximum intensity. To enable precise inter-spot alignment, each cropped region was registered to a consistent reference position within the subvolume coordinate system (11, 11, 5). All cropped spot images were voxel-wise averaged to generate an average 3D spot image (Fig. S2a). The background, defined as the mean intensity of the first axial slice, was subtracted from all voxel intensities.

One-dimensional intensity profiles were extracted through the centroid along $x$, $y$, and $z$ axes. The full width at half maximum (FWHM) was calculated using linear interpolation at half the profile's maximum intensity. FWHM values in pixels were converted to micrometers using voxel dimensions (0.220 µm lateral, 1 µm axial), resulting in 0.66 µm, 0.71 µm, and 2.06 µm for $x$, $y$, and $z$ respectively (Fig. S2b).

Presuming that the intensity profiles are gaussian, the lateral FWHM values were converted to a spatial standard deviation using $\sigma = \text{FWHM}/2.355$. The axial $\sigma_z$ includes an additional term for the variance of a uniform distribution for a 1-micron step over the 12 planes:

$$\sigma_z = \sqrt{\left(\frac{(1\ \mu\text{m}}{\sqrt{12}}\right)^2 + \left(\frac{\text{FWHM}_z}{2.355}\right)^2}. \tag{S1}$$

To additionally characterize the point spread function, we used PSFJ [9] to analyze images of 50 nm fluorescent beads with a lateral pixel size of 101 nm and a $z$-step of 250 nm. This yielded FWHM values of 0.44 µm, 0.49 µm, and 1.54 µm for $x$, $y$, and $z$, respectively (Fig. S2b).

The FWHM from averaged loci is larger in each dimension due to experimental factors such as motion blur over the 10-second temporal resolution, a 4x larger $z$-step, and the progressive accumulation of MS2 signal from polymerases elongating along the gene. Because these factors will contribute to the errors in the trajectories, we use the loci values to define the localization error for distance between two loci as

$$\sigma_\text{R} = \sqrt{2\sigma_x^2 + 2\sigma_y^2 + 2\sigma_z^2}. \tag{S2}$$

Based on this value ($\sigma_\text{R} = 1.43$ µm), we consider results for $R > \sigma_\text{R}$.

###### 2.2.7 Shuffling data as a control

To check whether the two allele fluctuation-distance profiles show a statistically significant effect, we shuffle the experimental data to mix two allele tracks from different nuclei. Briefly, for each locus position track, we first subtract the nucleus center coordinates. This gives us locus pair tracks in the reference frame of the nucleus. Next, we draw random pairs of nuclei $\{(p_1, p_2)_i\}_{i \in 1,\ldots,n}$, to create a shuffled data-set where the position of locus 1 is taken from nucleus $p_1$ and the position of locus 2 from nucleus 2. This way, we construct two-locus tracks where the labeled loci are from different nuclei, and hence their motion is expected to be uncorrelated. As expected, $\phi$ for the shuffled data look flat (Main Fig. 2h).

#### 2.3 ATP depletion in mESCs.

##### 2.3.1 Cell culture

New imaging data for the ATP depletion experiment have been acquired on a previously established cell line, based on E14Tg2a mouse embryonic stem cells, carrying LacO and TetO operator arrays separated by 150 kb [10]. Cells were cultured on gelatin-coated culture plates in Glasgow Minimum Essential Medium (GMEM) (Sigma-Aldrich, G5154) supplemented with 15% foetal calf serum (Eurobio Abcys), 1%L-Glutamine (Thermo Fisher Scientific, 25030024), 1% Sodium Pyruvate MEM (Thermo Fisher Scientific, 11360039), 1% MEM Non-Essential Amino Acids (Thermo Fisher Scientific, 11140035), 100 µM $\beta$-mercaptoethanol (Thermo Fisher Scientific, 31350010), 20U/ml leukemia inhibitory factor (Miltenyi Biotec, premium grade) in 8% CO2 with 2i inhibitors (1µM MEK inhibitor PDO35901 (Axon, 1408) and 3µM GSK3 inhibitor CHIR 99021 (Axon, 1386)) at 37°C. For live-cell imaging experiments, 106 cells were seeded on 35-mm glass-bottom dishes (Mattek, P35G1.5-14-C) coated with 2 µg ml-1 laminin (Sigma-Aldrich, L2020) in PBS at 37°C overnight and cultured for 24 h in Fluorobrite Dulbecco's Modified Eagle Medium (DMEM) (Gibco, A1896701) supplemented with 15% foetal calf serum (Eurobio Abcys), 1% L-Glutamine (Thermo Fisher Scientific, 25030024), 1% Sodium Pyruvate MEM (Thermo Fisher Scientific, 11360039), 1% MEM Non-Essential Amino Acids (Thermo Fisher Scientific, 11140035), 100µM $\beta$-mercaptoethanol (Thermo Fisher Scientific, 31350010), 20U/ml leukemia inhibitory factor (Miltenyi Biotec, premium grade) and 2i inhibitors. For ATP depletion, the medium was replaced with Fluorobrite medium containing 1 µM rotenone (Tocris-Bioscience) and 6 mM 2-Deoxy-D-glucose (Tocris-Bioscience) 1 hour prior to the imaging session [11].

##### 2.3.2 Microscope setup and live-cell imaging acquisition

Cells were imaged with a Nikon Eclipse Ti2 inverted widefield microscope equipped with a Total Internal Reflection Microscopy iLAS2 module (Gataca systems) for azimuthal illumination [12], a Perfect Focus System (Nikon) and motorized Z-Piezo stage (ASI) using a CFI APO TIRF 100×, 1.49NA oil immersion objective (Nikon). The microscope was operated in oblique illumination mode with a nominal incidence angle of 54°. Excitation sources were a 473 nm and a 552 nm Omicron laserbench laser. Cells were maintained at 37 °C and 8% CO2 using an enclosed microscope environmental control setup (Cube) and CO2 control (Brick) from Life science instruments. The microscope was equipped with two dual back-illuminated 95% quantum efficiency scientific CMOS cameras (Orca-fusion-BT, Hamamatsu) with a pixel size of 6.5 µm. A dichroic long-pass (550, Chroma) was added to separate red from green to the two cameras. Furthermore, two bandpass (514/44, Semrock, 595/65, Chroma, respectively) filters were added to the green and red channel.

The microscope was controlled using NIS-Elements software (Nikon, version 5.42.07).

Laser power was set to 2% of the maximum power for the 473 nm and 3% 552 nm lasers which approximately corresponds to powers of 695 µW and 225 µW at the objective. The field of view was binned 2 × 2 in the $x$ and $y$ directions which resulted in an effective pixel size of 130 nm. Images were acquired for 5 minutes at 0.5 Hz (2 s between frames) with an exposure time of 100 ms per channel and a $z$-step size of 0.3 µm for 9 stacks.

##### 2.3.3 Image processing and construction of LacO-TetO distance tracks

Images were saved in the NIS-Elements format (.nd2) and processed in Python using the nd2 library available on GitHub. We employed the same imaging analysis pipeline used in [4] available on GitHub at [13].

To remove uncorrected artifacts from the imaging pipeline, we further refined the distance tracks by manually inspecting frames where the square displacements between successive frames (2 s) were greater than 200 nm$^2$, corresponding to approximately two times the mean-square displacement measured at 2 s intervals in [4]. This way, we filtered out a few tracks with replicated chromosomes, where the switching of detection between two sister chromatids from one frame to the next caused large displacements and some mislocalizations (Fig. S3). This control removed approximately 10% of the tracks from the initial processed dataset.

To estimate cell diameters, we used the 2D segmentations of the imaged cells and computed the lengths of their corresponding major and minor principal axes using the major_axis_length and minor_axis_length properties of the regionprops function from the scikit-image Python package [14]. These properties return the lengths of the axes of an ellipse that approximates the segmented cell shape. The mean major and minor axis lengths were 15 ± 2 µm and 11 ± 2 µm, respectively.

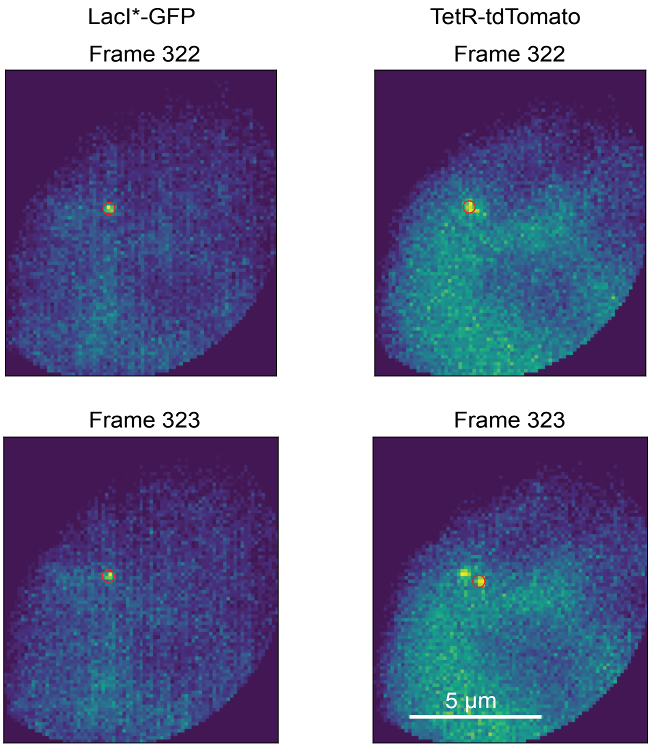

Figure S3: **Spot localization in mESC experiments.** Example of spot localizations in successive frames for the two operator arrays, illustrating an event that results in a large displacement in the vector between the operators. This artifact is caused by switching of the TetR-tdTomato localization between two replicated spots from one frame to the next.

### 3 Simulation models

#### 3.1 Rouse model without SCFs

As a null model to compare our SCFs simulations to, we consider a linear 3D Rouse polymer confined by a harmonic potential to a sphere of diameter $D$ centered at the origin, as a minimal model of the confinement by the cell nucleus. The polymer consists of $N$ monomers, whose over-damped motion is described by:

$$d\mathbf{r}_i = -\frac{k}{\zeta}(2\mathbf{r}_i - \mathbf{r}_{i-1} - \mathbf{r}_{i+1})dt + \frac{F_{\text{conf}}(|\mathbf{r}_i|, D)}{\zeta}dt + d\boldsymbol{\xi}_i(\{\mathbf{r}_j(t)\}_j, t) \tag{S3}$$

for all monomers $1 < i < N$. For the ends of the polymer, corresponding to monomers $i = 1$ and $i = N$, only one spring term is present, equivalent to defining $\mathbf{r}_0 = \mathbf{r}_1$ and $\mathbf{r}_{N+1} = \mathbf{r}_N$.

The spring stiffness $k = k_\text{B}T/l_0^2$ is set by the effective thermal energy (effective temperature $T$ times Boltzmann constant $k_\text{B}$) and the equilibrium length of the spring $l_0$.

The uncorrelated noise in the Rouse model $d\boldsymbol{\xi}_i(\{\mathbf{r}_j(t)\}_j, t)$ is given by i.i.d. white noise increments, with zero mean and covariance

$$\langle d\xi_{i,\alpha}(t) d\xi_{j,\alpha'}(t')\rangle = \frac{2k_\text{B}T\,dt}{\zeta}\delta_{i,j}\delta_{\alpha,\alpha'}\delta(t-t') =: \sigma^2 dt\ \delta_{i,j}\delta_{\alpha,\alpha'}\delta(t-t'), \tag{S4}$$

where $\alpha$ and $\alpha'$ are vector indices. In the next section, we will introduce spatially correlated noise increments.

To compare simulations with fly embryo data [2], we fix simulation units based on experimental data. We coarse-grain a 4000 kb chromosomal section into $N = 800$ beads, each corresponding to 5 kb. $l_0$ is set by matching the mean extension of 391 nm for the shortest considered genomic section of length 58 kb, or $n \approx 12$ beads;

$$l_0 = \sqrt{\frac{\langle R(n)^2\rangle}{n}} \approx 113 \text{ nm}. \tag{S5}$$

While the radius of gyration of an unconfined Rouse chain is given by $R_g = \sqrt{\langle R^2\rangle} \sim s^{1/2}$, experimentally a shallower scaling is observed which is approximately matched by our confined Rouse simulation (Fig. S4a,b). Unless otherwise stated, we therefore present simulation results in the presence of a confinement potential. Importantly, however, we find that in simulations without spatial confinement, the scalings of $\phi(R, \Delta t)$ remain unchanged (SI Appendix Fig. S8).

The diameter of the confinement potential is selected as $D = 2600$ nm, based on the maximal extensions observed in experiments. The confinement force is defined as

$$F_{\text{conf}}(|\mathbf{r}|, D) = \begin{cases} 0; \text{ if } |\mathbf{r}| < D/2 \\ -k_{\text{conf}}(|\mathbf{r}| - D/2))\hat{\mathbf{r}}; \text{ else,} \end{cases} \tag{S6}$$

where $\hat{\mathbf{r}}$ is a unit vector along $\mathbf{r}$, and $k_{\text{conf}}$ is the confinement potential strength. To ensure that fluctuations far outside the confinement are rare, we set $k_{\text{conf}} = 2k_{\text{B}}T/w_{\text{conf}}^2$, where $w_{\text{conf}} = 400$ nm defines the effective width of the potential.

Finally, the time-scale of the simulations is determined by setting the magnitude of the frictional force, $\zeta$. We set this by matching the single-locus diffusivity $\Gamma_1 = \langle \Delta r^2 \rangle \Delta t^{-1/2}$ at the imaging time-interval $\Delta t = 28$ s [15]:

$$\zeta = \frac{2k_{\text{B}}Tl_0^2}{\pi\,\Gamma_1^2} \tag{S7}$$

We simulate the system using a custom code in Julia (available online, see data availability). We first initialize the polymer as a random walk within the confinement sphere, and let it relax within the confinement before starting sampling. We integrate Eq. (S3) numerically, using $\mathbf{r}_i(t+dt) = \mathbf{r}_i(t) + d\mathbf{r}_i(t)$ with time-step $dt = 0.005\zeta/k$. For uncorrelated Rouse simulations, the noise for each monomer at each time step is sampled from a normal distribution with standard deviation $\sigma$.

All parameters of the model were constrained using experimental data (Table S1), and both $\langle \Delta r^2 \rangle$ and $\langle R \rangle$ match between simulation and experiment (Fig. S4b). Hence, the absence of a strong $s$-dependence of $\Gamma_2$ in the simulations suggests that COM correlations between locus motions cannot explain the experimentally observed $\Gamma_2(s)$ curve (Main Fig. 1b).

Table S1: Parameters of the confined Rouse model without SCFs. $\Gamma_1$ is inferred from single-locus MSDs in fly embryos.

| Parameter | Symbol | Value |
|---|---|---|
| Number of monomers | $N$ | 800 |
| Coarse-graining resolution | — | 5 kb/bead |
| Chromosomal section length | — | 4000 kb |
| Equilibrium spring length | $l_0$ | 113 nm |
| Effective thermal energy | $k_{\text{B}}T$ | 4.114 pN nm |
| Spring stiffness | $k$ | $k_{\text{B}}T/l_0^2$ |
| Confinement diameter | $D$ | 2600 nm |
| Confinement width | $w_{\text{conf}}$ | 400 nm |
| Confinement stiffness | $k_{\text{conf}}$ | $2\,k_{\text{B}}T/w_{\text{conf}}^2$ |
| Single locus diffusivity | $\Gamma_1$ | 18900 $\text{nm}^2\text{s}^{-1/2}$ |
| Friction coefficient | $\zeta$ | $2\,k_{\text{B}}T\,l_0^2/(\pi\,\Gamma_1^2)$ |
| Noise amplitude | $\sigma^2$ | $2\,k_{\text{B}}T/\zeta$ |
| Integration time step | $dt$ | $0.005\,\zeta/k$ |
| Imaging time interval | $\Delta t$ | 28 s |

##### 3.2 Rouse model in correlated flow field

To model SCFs arising from active flows within the nucleoplasm, we include a minimal implementation of a random flow field $\mathbf{u}(\mathbf{x}, t)$ in our simulations. At each simulation

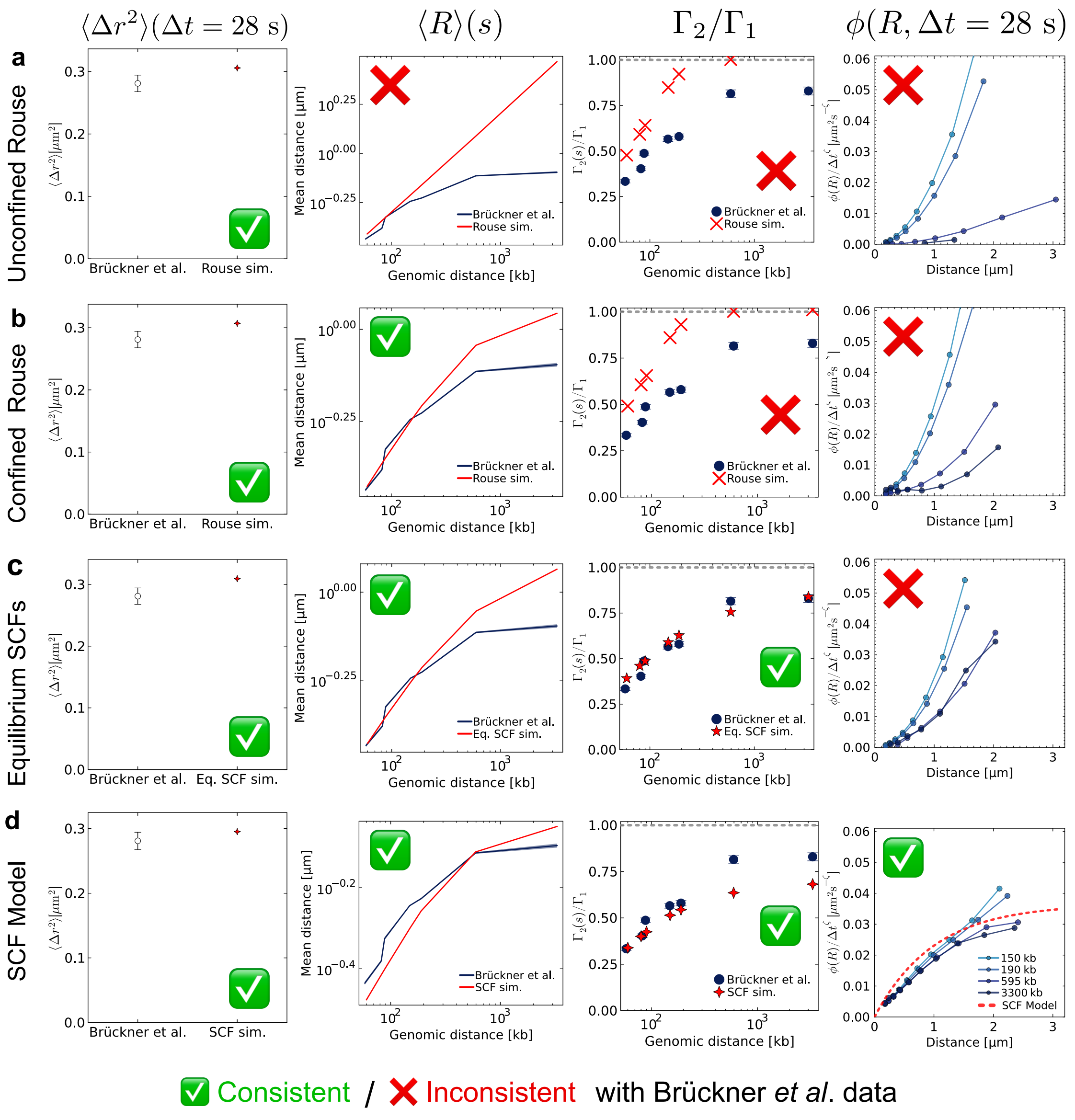

Figure S4: **Comparison of 3D simulations. (a)** $\langle \Delta r^2 \rangle$, $\langle R \rangle(s)$, $\Gamma_2/\Gamma_1$, and $\phi(R, \Delta t = 28\ \text{s})$ plots for 3D Rouse simulations without confinement. Green ticks indicate consistency with experimental data from [2], red crosses indicate qualitative differences compared to experimental data. **(b)** Similar plots for 3D Rouse model confined to a sphere. The mean distances between loci now show the correct scaling with $s$. **(c)** Similar plots for 3D simulations with spatially correlated equilibrium white noise. Although the model captures the $\Gamma_2/\Gamma_1$ curves, it does not show the experimentally observed $\phi$ scaling at $\Delta t = 28$ s. **(d)** Similar plots for 3D SCF simulations with spatially correlated active flows. The model is qualitatively consistent with all experimental statistics. Results in 2D are equivalent (Fig. S6).

step, we update the positions of all monomers according to

$$dr_{i,\alpha} = \underbrace{-\frac{F_{i,\alpha}}{\zeta}dt + \sigma dW_{i,\alpha}}_{\text{Uncorrelated Rouse model}} + \underbrace{u_\alpha(\mathbf{r_i})dt}_{\text{Flow}}. \tag{S8}$$

To construct the random spatially correlated flow-field, we first set up the following arrays:

1. We define a spatial grid with lattice spacing $dx = \lambda_{\text{flow}}/4.5$ within the volume of confinement. We hence have a set of evenly spaced grid points $\{\mathbf{x}_i\}_i$.
2. Given the correlation length $\lambda_{\text{flow}}$, we calculate the filter $I(\mathbf{x}_i) = I_0 e^{-|\mathbf{x}|_i/\lambda_{\text{flow}}}$ on the lattice.
3. We precompute the Fourier transform of $\hat{I}(\mathbf{k})$.
4. We initialize arrays $\mathbf{u}(\mathbf{r}_i)$ on the lattice and $\hat{\mathbf{u}}(\mathbf{k})$ and $\hat{\mathbf{u}}_{\text{proj}}(\mathbf{k})$ in reciprocal space.

At each time-step

1. We draw a white noise field $\boldsymbol{\xi}(\mathbf{x_i})$ on the lattice.
2. We Fourier transform the noise field; $\hat{\boldsymbol{\xi}}(\mathbf{k}) = \text{fft}(\boldsymbol{\xi}(\mathbf{x}_i))$, where $\mathbf{k}$ is a wave-vector.
3. We calculate $\hat{I} \times \hat{\boldsymbol{\xi}}$, corresponding to the Fourier transform of the convolution of the noise field with the filter $I$. This ensures that the flow field has the desired correlation length.
4. We update $\hat{\mathbf{u}}(\mathbf{k})$ as an Ornstein-Uhlenbeck process with decay time $T_{\text{flow}}$: $\hat{\mathbf{u}}(\mathbf{k}, t + dt) = e^{-dt/T_{\text{flow}}}\hat{\mathbf{u}}(\mathbf{k}, t) + \sqrt{1 - e^{-2dt/T_{\text{flow}}}}\hat{I} \times \hat{\boldsymbol{\xi}}$.
5. We project $\hat{\mathbf{u}}$ to a divergence-free sub-space by defining $\hat{\mathbf{u}}_{\text{proj}}(\mathbf{k}) = (1 - \mathbf{k}\mathbf{k}^T/k^2) \times \hat{\mathbf{u}}(\mathbf{k})$.
6. We set $\hat{\mathbf{u}}_{\text{proj}}(\mathbf{0}) = 0$. This removes the constant term of the flow field. Physically, we assume the active flow conserves momentum.
7. Finally, we define $\mathbf{u} = \text{ifft}(\hat{\mathbf{u}}_{\text{proj}})$ on the lattice.
8. We construct an interpolator that allows us to sample $\mathbf{u}(\mathbf{r}_i)$ for an off-lattice monomer position $\mathbf{r}_i$.

This process ensures that $\mathbf{u}$ represents a divergence-free flow field with zero-momentum, with correlations decaying across lengths $\lambda_{\text{flow}}$ and time $T_{\text{flow}}$. On time-scales $t \gg \tau$, the relative contribution of the flow field to the diffusion of a tracer particle is characterized by the dimensionless parameter

$$f_{\text{flow}} = \frac{\langle u^2 \rangle \tau \zeta}{2k_{\text{B}}T}. \tag{S9}$$

To set the parameters of the model, we proceed as follows. We start with the same parameters $\zeta,\ k, k_{\text{B}}T, D, N$ as in the Rouse model. Based on experimentally measured fluctuation-distance profiles $\phi(R)$ (Fig. S7, Table S4), we vary $\lambda_{\text{flow}}$ in the range 300-600 nm. We vary $T_{\text{flow}} \in [\Delta t/2, 2\Delta t]$; the results are relatively insensitive to the precise value within this range (Fig. S5). Given these parameters, we find an initial value for $f_{\text{flow}}$ such that the model matches the experimental $\Gamma_2(s)/\Gamma_1$ curve. This sets $\langle u^2 \rangle$ and $I_0 \propto \sqrt{\langle u^2 \rangle}$. The addition of the flow field speeds up locus motion relative to the Rouse model. To recalibrate the time-scale of the simulations to match experimental single-locus diffusivities, we rescale the friction coefficient $\zeta \to \zeta/w$ by a dimensionless factor $w = \langle \Delta r^2 \rangle_{\text{sim}} / \langle \Delta r^2 \rangle_{\text{exp}}$. To keep $f_{\text{flow}}$ constant, we also scale $\langle u^2 \rangle \to \langle u^2 \rangle w$. Given these rescalings, the single-locus diffusivities match experiments (Fig. S4).

We note that the parameters $\langle u^2 \rangle$ and $\lambda_{\text{flow}}$ are not individually uniquely constrained: Fig. S5 shows that a larger flow magnitude can be partially compensated by increasing $\lambda_{\text{flow}}$. However, using $\lambda_{\text{flow}}$ and $\langle u^2 \rangle$ as fitted parameters, the model predicts the full distance-dependent and time-dependent structure of $\phi(R, \Delta t)$ as well as $\Gamma_2(s)$ without further adjustment (Main Fig. 2).

Since 2D and 3D simulations of the model give consistent results (Fig. S6), we minimized computational cost by conducting flow simulations in 2D using a custom Julia script. This significantly reduces computational time since the number of operations in the Fourier transforms in the simulations scales with $L^d$ for a lattice with $L$ points in $d$ dimensions. Unless otherwise stated, we display 2D results.

Table S2: Additional parameters of the active flow field model. All other parameters are identical to the Rouse model (Table S1).

| Parameter | Symbol | Value |
|---|---|---|
| Flow correlation length | $\lambda_{\text{flow}}$ | [300, 600] nm |
| Flow decay time | $T_{\text{flow}}$ | $[\Delta t/2,\ 2\Delta t]$ |
| Relative flow field magnitude | $f_{\text{flow}}$ | set by matching $\Gamma_2/\Gamma_1$ |
| Friction rescaling factor | $w$ | $\langle \Delta r^2 \rangle_{\text{sim}} / \langle \Delta r^2 \rangle_{\text{exp}}$ |
| Rescaled friction coefficient | $\tilde{\zeta}$ | $\zeta/w$ |
| Flow field magnitude | $\langle u^2 \rangle$ | $2k_{\text{B}}T f_{\text{flow}}/(\tau\tilde{\zeta})$ |
| Filter normalization | $I_0$ | $\propto \sqrt{\langle u^2 \rangle}$ |
| Lattice spacing | $dx$ | $\lambda_{\text{flow}}/4.5$ |

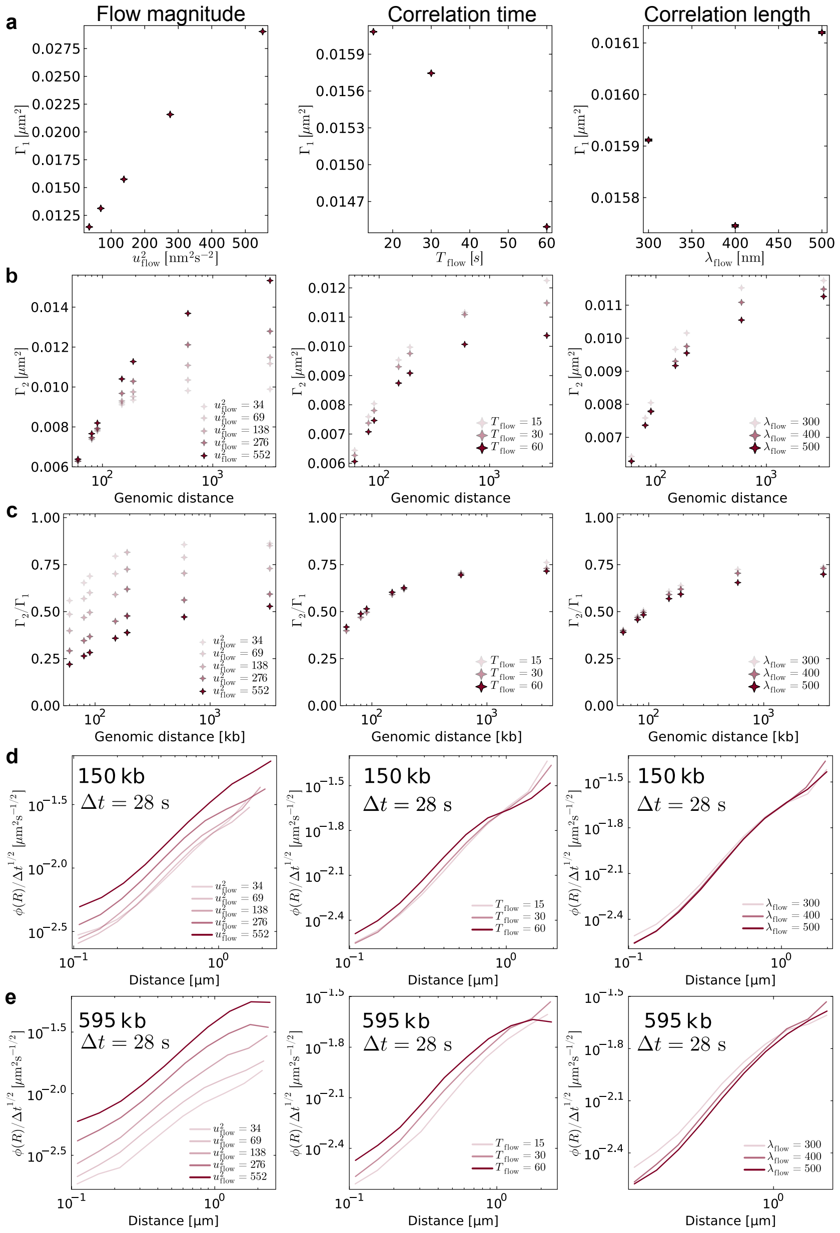

Figure S5: **Effects of parameters on SCF simulation results.** Table showing effects of varying average flow magnitude $u_{\text{flow}}$, flow correlation time $T_{\text{flow}}$, or flow correlation length $\lambda_{\text{flow}}$ on... **(a)** single-locus diffusivity. Stronger flows increase diffusion. A larger correlation time slightly decreases diffusion, whereas the correlation length has a negligible effect. **(b)** two-locus diffusivity. Stronger flows increase two-locus diffusivity, especially for larger genomic separations. Increasing flow correlation time or length slightly decreases two locus diffusivities, consistent with locus motion becoming more correlated. **(c)** the ratio $\Gamma_2/\Gamma_1$. Increasing flow magnitudes increase $\Gamma_1$ faster than $\Gamma_2$, resulting in a larger discrepancy between $\Gamma_2$ and $\Gamma_1$. $T_{\text{flow}}$ and $\lambda_{\text{flow}}$ have small effects on the ratio. **(d)** Fluctuation-distance profiles for $s = 150$ kb. Larger flow magnitudes increase correlated motion, and hence shift the magnitude of $\phi$. $T_{\text{flow}}$ and $\lambda_{\text{flow}}$ have subtle effects on the shapes of the profiles. **(e)** $\phi$ for $s = 595$ kb. The effects of all parameters are stronger for larger $s$.

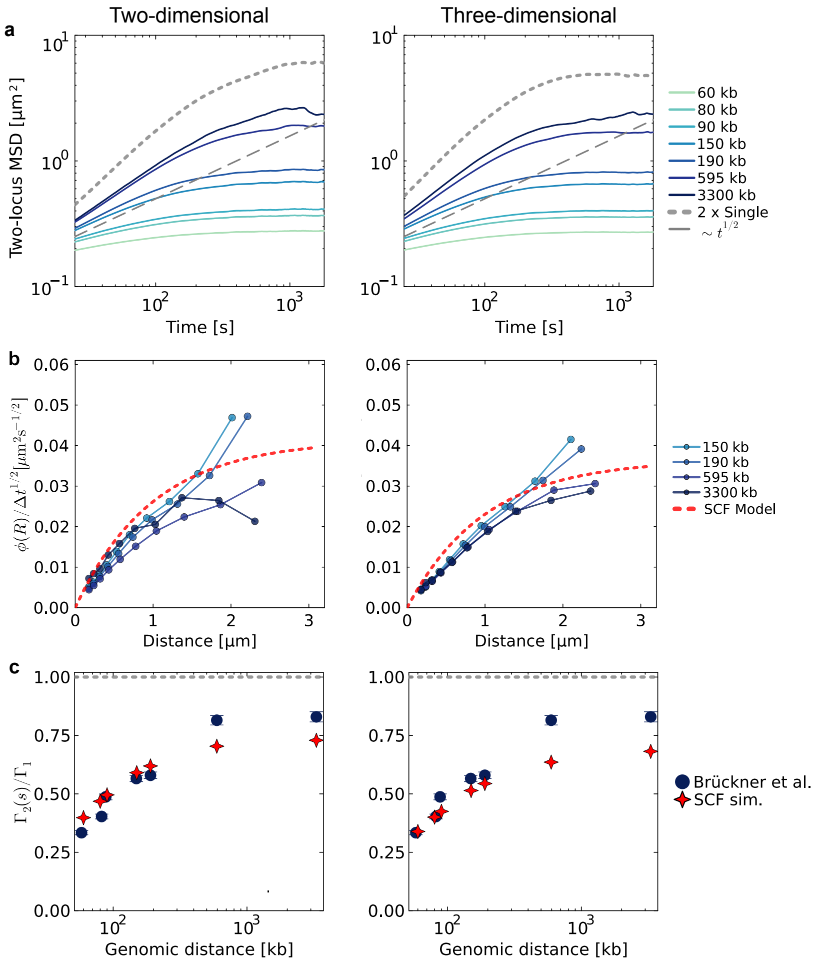

Figure S6: **Flow simulations give similar results in two or three dimensions (a)** MSD curves for flow simulations in two or three dimensions. The curves are scaled by $3/d$ to ensure comparable magnitudes. Same parameters ($u_{\text{flow}} = 11.8$ µm s$^{-1}$, $T_{\text{flow}} = 30$ s, $\lambda_{\text{flow}} = 400$ nm) used in both simulations. **(b)** Similar comparison of $\phi(R)$ curves, using $\Delta t = 28$ s. To compare magnitudes, curves are scaled by $1/d$. **(c)** Similar comparison of $\Gamma_2(s)$ curves. Since both $\Gamma_1$ and $\Gamma_2$ are expected to scale with $d$, no normalization of the curves is required.

##### 3.3 Rouse model with spatially correlated equilibrium white noise

The Rouse model assumes spatially and temporally uncorrelated noise on each monomer, meaning that the noise is $\delta$-correlated in both time and space. To extend the Rouse model to include spatially correlated noise while maintaining thermodynamic equilibrium, we develop a model in which the noise driving each monomer is temporally white (i.e., $\delta$-correlated in time) but spatially correlated between monomers. For the model to be at equilibrium, the noise must satisfy the fluctuation-dissipation relation, so that the system relaxes to the Boltzmann distribution $P_{\mathrm{eq}} \propto e^{-U/k_{\mathrm{B}}T}$ at long times, where $U$ is the total potential energy of the polymer configuration. This is in contrast to the active flow model (Section 3.2), where temporally correlated flows break detailed balance and drive the system out of equilibrium.

Because the noise correlations depend on the instantaneous distances between monomers, $C_{\mathrm{cor}}(|\mathbf{r}_i - \mathbf{r}_j|)$, the noise is state-dependent: its amplitude depends on the current state of the system. Such state-dependent noise gives rise to a noise-induced drift term in the Langevin equation [16]. This drift is required for thermodynamic consistency: without it, the Fokker-Planck equation associated with the Langevin dynamics does not have the Boltzmann distribution as its steady-state solution. The specific form of the drift depends on the stochastic calculus convention used to interpret the multiplicative noise (e.g., Itô vs. Stratonovich). Here, we follow the Itô convention and apply the multi-component framework of ref. [16] to our polymer system. We provide a brief derivation for completeness.

Let $U(\{\mathbf{r}_k\}_{k\in\{1,\dots,N\}})$ be the potential energy of a polymer configuration, including contributions from both the spring forces between beads and the confinement potential. We look for forces $F$ such that the process for each dimension $\alpha$ is defined by

$$dr_{i,\alpha} = F_{i,\alpha}(\{\mathbf{r}_k\})dt + \sigma_{i,j}(\{\mathbf{r}_k\})dW_{j,\alpha} \tag{S10}$$

gives rise to the Boltzmann equilibrium distribution (i.e., the same equilibrium distribution as in the uncorrelated noise case); $P_{\mathrm{eq}}(\{\mathbf{r}_k\}) = e^{-U(\{\mathbf{r}_k\})/k_{\mathrm{B}}T}/Z$, where $Z$ is the partition function. Here, repeated indices indicate summation, and each $dW_{j,\alpha}$ is a standard Wiener process. Correlations are implemented by defining the matrix $\sigma$ such that $\sum_l \sigma_{i,l}\sigma_{j,l} = C_{\mathrm{cor}}(|\mathbf{r}_i - \mathbf{r}_j|)$. In the uncorrelated case, we would have $\sigma_{i,j} = \sqrt{\frac{2k_{\mathrm{B}}T}{\zeta}}\delta_{i,j}$.

By writing down the Fokker-Planck equation, and by assuming that at steady-state, the probability current of the process must be zero, we find

$$0 = F_{i,\alpha}(\{\mathbf{r}_k\})P_{\mathrm{eq}}(\{\mathbf{r}_k\}) - \sum \frac{\partial}{\partial r_{j,\alpha}}[D_{i,j}(\{\mathbf{r}_k\})P_{\mathrm{eq}}(\{\mathbf{r}_k\})] \tag{S11}$$

where $\alpha$ indicates the dimension index of a vector, and

$$D_{i,j}(\{\mathbf{r}_k\}) = \frac{C_{\mathrm{cor}}(|\mathbf{r}_i - \mathbf{r}_j|)}{2} \tag{S12}$$

is the diffusion matrix that encodes the spatial noise correlations.

Solving for $F_{i,\alpha}$ yields

$$F_{i,\alpha}(\{\mathbf{r}_j\}) = \sum_j \frac{\partial D_{i,j}}{\partial r_{j,\alpha}} - \frac{D_{i,j}}{k_\mathrm{B}T}\frac{\partial U}{\partial r_{j,\alpha}}. \tag{S13}$$

The first term is the noise-induced drift arising from the state-dependence of the diffusion matrix. In the case of uncorrelated motion, $D_{i,j} = \delta_{i,j}k_\mathrm{B}T/\zeta$, the drift vanishes and we recover the standard overdamped Langevin equation:

$$F_{i,\alpha}(\{\mathbf{r}_j\}) = -\frac{1}{\zeta}\frac{\partial U}{\partial r_{i,\alpha}}. \tag{S14}$$

To summarize, noise correlations affect the forces in two ways. First, forces exerted on bead $j$ also affect the motion of bead $i$ via the off-diagonal elements of the diffusion matrix. Second, the state-dependence of the noise introduces a drift term proportional to the spatial derivatives of the diffusion tensor, which is required for the system to relax to the correct equilibrium distribution.

To include spatially correlated fluctuations that are consistent with equilibrium white noise in our simulation model, we assume that noise correlations depend on the instantaneous distance between loci. In short, we define $\boldsymbol{\xi}_i(\{\mathbf{r}_j(t)\}_j, t)$ in Eq. (S3) such that

$$\langle d\xi_{i,\alpha}(\{\mathbf{r}_k(t)\}_k, t) d\xi_{j,\alpha'}(\{\mathbf{r}_l(t')\}_l, t')\rangle = 2k_\mathrm{B}T dt \delta_{\alpha,\alpha'}\delta(t-t') \times \tag{S15}$$

$$\left(\frac{f_\mathrm{cor}e^{-|\mathbf{r}_i-\mathbf{r}_j|/\lambda_\mathrm{cor}}}{\zeta'} + \frac{\delta_{i,j}(1-f_\mathrm{cor})}{\zeta'}\right). \tag{S16}$$

Here, $\zeta'$ is a rescaled friction constant, as discussed below. The two free parameters are $\lambda_\mathrm{cor}$, the length scale at which the noise correlations decay, and $f_\mathrm{cor}$, which scales the noise correlations with respect to independent noise. We keep the parameters $k, k_BT, D$ the same as in the Rouse simulations (Eq. (S7), Table S1), but adjust $\zeta' \neq \zeta$ such that the single-locus diffusivity $\Gamma_1$ still matches experiments.

Following the theoretical analysis presented in Section 3.3, at each time-step we calculate the following arrays:

1. The sum of the spring and confinement forces acting on each bead $i$; $\mathbf{F}_i$.
2. The diffusion matrix $D_{i,j} = k_BTf_\mathrm{cor}e^{-|\mathbf{r}_i-\mathbf{r}_j|/\lambda_\mathrm{cor}}/\zeta'$.
3. The derivatives of the diffusion matrix; $\partial D_{i,j}/\partial r_{j,\alpha} = D_{i,j}/\lambda_\mathrm{cor} \times (r_{i,\alpha} - r_{j,\alpha})/|\mathbf{r}_i - \mathbf{r}_j|$.
4. We perform a Cholesky decomposition to find $L$ such that $L^TL = 2D$.
5. We sample an arrays of noise: $\mathbf{W}$ of size (N,3).

Given these arrays, the position of monomer $i$ is updated by

$$dr_{i,\alpha} = \underbrace{-\frac{1-f_{\text{cor}}}{\zeta'}F_{i,\alpha}dt + \sqrt{(1-f_{\text{cor}})}\sigma' dW_{i,\alpha}}_{\text{Uncorrelated Rouse model}} + \underbrace{\sum_j \left[\left(\frac{\partial D_{i,j}}{\partial r_{j,\alpha}} - \frac{D_{i,j}}{k_B T}F_{j,\alpha}\right)dt + L_{i,j}dW_{j,\alpha}\right]}_{\text{Equilibrium SCF terms}}. \tag{S17}$$

Here $\sigma' = 2k_\text{B}Tdt/\zeta'$. Note that in the limit $f_{\text{cor}} \to 0$, $D = L = 0$, and we recover the uncorrelated Rouse simulation.

In general, if we set $\zeta'$ to the same value $\zeta$ as in uncorrelated Rouse simulations, the time-scales of the simulations would change. To ensure the deterministic displacements in a given time interval are of the same magnitude, we rescale the friction such that the magnitude of deterministic forces acting on each bead remains constant:

$$\left\langle \frac{|\mathbf{F}_i|}{\zeta} \right\rangle = \left\langle \frac{|D_{ij}\mathbf{F}_j|}{k_B T} \right\rangle, \tag{S18}$$

where the averages are taken over all monomers $i$ in different polymer configurations from Rouse simulations. Since the systems have the same equilibrium distributions, these averages are the same in the correlated models. We hence find

$$\zeta'(\lambda_{\text{cor}}) = \zeta \left\langle |\sum_j e^{-|\mathbf{r}_i - \mathbf{r}_j|/\lambda_{\text{cor}}} \mathbf{F}_j| \right\rangle / \langle |\mathbf{F}_i| \rangle. \tag{S19}$$

We find that this rescaling ensures that $\Gamma_1 = \langle \Delta r^2 \rangle \Delta t^{-1/2}$ remains roughly constant as we vary $f_{\text{cor}}$ and $\lambda_{\text{cor}}$ (Fig. S4).

After fixing $k$ based on the radius of gyration data, and $\zeta'$ based on $\lambda_{\text{cor}}$ and $\zeta$, the only free variables in the correlated model are $f_{\text{cor}}$ and $\lambda_{\text{cor}}$.

The equilibrium simulations were conducted in 3D using a custom Julia code, making use of CUDA.jl [17, 18]. Simulations were performed on NVIDIA A100 GPUs.

Table S3: Additional parameters of the equilibrium spatially correlated fluctuations (SCF) model. All other parameters are identical to the Rouse model (Table S1).

| Parameter | Symbol | Value |
|---|---|---|
| Noise correlation length | $\lambda_{\text{cor}}$ | free parameter |
| Noise correlation fraction | $f_{\text{cor}}$ | free parameter |
| Rescaled friction | $\zeta'$ | $\zeta \left\langle \left\| \sum_j e^{-\|\mathbf{r}_i - \mathbf{r}_j\|/\lambda_{\text{cor}}} \mathbf{F}_j \right\| \right\rangle / \langle \|\mathbf{F}_i\| \rangle$ |
| Rescaled noise amplitude | $\sigma'^2$ | $2\, k_\text{B} T\, dt/\zeta'$ |

### 4 Supplementary theory

#### 4.1 Decomposing $\Gamma_2$

To derive Eq. 1, we start by writing [19]

$$\mathrm{MSD}_2(\Delta t) = \langle \Delta R^2 \rangle = \langle (\mathbf{R}(t+\Delta t) - \mathbf{R}(t))^2 \rangle = 2\Gamma_2 \Delta t^{\beta}. \tag{S20}$$

By solving for $\Gamma_2$ and substituting $\mathbf{R} = \mathbf{r}_i - \mathbf{r}_j$, we find

$$\begin{aligned}\Gamma_2 &= 1/2\ \Delta t^{-\beta} \langle (\mathbf{r}_i(t+\Delta t) - \mathbf{r}_j(t+\Delta t) - \mathbf{r}_i(t) + \mathbf{r}_j(t))^2 \rangle & \text{(S21)}\\ &= 1/2\ \Delta t^{-\beta} \langle (\Delta \mathbf{r}_i - \Delta \mathbf{r}_j)^2 \rangle. & \text{(S22)}\end{aligned}$$

Assuming that single locus dynamics are identical, we have $\langle \Delta \mathbf{r}_i^2 \rangle = \langle \Delta \mathbf{r}_j^2 \rangle = \Gamma_1 \Delta t^{\beta}$. Expanding the square in Eq. (S22), we hence find the result

$$\Gamma_2 = \Gamma_1 - \langle \Delta \mathbf{r}_i \cdot \Delta \mathbf{r}_j \rangle\ \Delta t^{-\beta}. \tag{S23}$$

#### 4.2 Theoretical predictions for $\phi$

##### 4.2.1 Derivation of $\phi$ from the Rouse model

To derive the scaling of the fluctuation-distance profile $\phi$, we start by analyzing the scaling behavior of $\langle \Delta R^2 | R \rangle$ for a simple Rouse chain. We consider the relaxation of Rouse modes $X_p$, each of which is described by an independent Ornstein-Uhlenbeck process [20, 21]. Intuitively, constraining $R(0)$ constrains the initial mode amplitudes $X_p(0)$. This introduces an $R$-dependence on $\langle \Delta R^2 | R \rangle$.

For simplicity, we consider a 1D system with $\mathbf{r}_i = r_i$ and $\mathbf{R} = R$. The results can be extended for $d$ dimensions by treating each dimension independently. We consider normal modes $\phi_p(s) = \cos(p\pi s/N)$ with amplitudes $X_p$, and write

$$r_i = X_0 + 2 \sum_{p=1}^{N} X_p \phi_p(i). \tag{S24}$$

The end-to-end length of the chain is described by

$$R = r_N - r_0 = -4 \sum_{p \text{ odd}} X_p. \tag{S25}$$

Each mode evolves independently according to

$$\zeta_p dX_p = -k_p X_p dt + \sigma_p dW_p. \tag{S26}$$

Here, for $p \ll N$, $k_p = \frac{\pi^2 k p^2}{N}$, and $\zeta_p = N\zeta$. $dW_p$ is a Wiener process with zero mean and variance $dt$, and the noise amplitude is $\sigma_p = \sqrt{2\zeta_p k_B T}$.

The time covariance of each mode is

$$\langle X_p(t)X_p(0)\rangle = \langle X_p^2\rangle e^{-t/\tau_p}, \tag{S27}$$

where $\tau_p = \zeta_p/k_p = \tau_1/p^2$. We can also consider a conditional average

$$\langle X_p(t)X_p(0)|R(0)\rangle = \langle X_p^2|R\rangle e^{-t/\tau_p}. \tag{S28}$$

Here, we use $\langle X_p^2|R\rangle := \langle X_p(t)^2|R(t)\rangle$ to indicate a conditional average where both quantities are evaluated at the same time-point. Using these expressions, we can also find the expected change in amplitude over a period $\Delta t$:

$$\langle \Delta X_p^2\rangle := \langle (X_p(\Delta t) - X_p(0))^2\rangle = 2\langle X_p^2\rangle(1 - e^{-\Delta t/\tau_p}). \tag{S29}$$

If we take a conditional average, given $R(t=0)$, we have

$$\langle \Delta X_p^2|R\rangle := \langle (X_p(\Delta t) - X_p(0))^2|R(0)\rangle = \langle X_p^2|R\rangle(1 - 2e^{-\Delta t/\tau_p}) + \langle X_p(\Delta t)^2|R(0)\rangle. \tag{S30}$$

We next use these results to find an expression for $\langle \Delta R^2|R\rangle$, for $\Delta t \ll \tau_1$. Since all modes evolve independently and have zero mean, we may write

$$\langle \Delta R^2\rangle = 16 \sum_{p \text{ odd}} \langle \Delta X_p^2\rangle. \tag{S31}$$

For the conditional average, by contrast, cross terms must be included:

$$\langle \Delta R^2|R\rangle = \underbrace{16 \sum_{p \text{ odd}} \langle \Delta X_p^2|R\rangle}_{\text{squared terms}} + \underbrace{16 \sum_{p\neq q \text{ odd}} \langle \Delta X_p|R\rangle\langle \Delta X_q|R\rangle}_{\text{cross terms}}. \tag{S32}$$

Here, we used that the modes $q \neq p$ are relaxing independently, but the conditional averages $\langle \Delta X_p|R\rangle$ are expected to be non-zero.

We first consider the scaling of the cross-terms, which include factors

$$\langle \Delta X_p|R\rangle = \langle X_p(\Delta t)|R(0)\rangle - \langle X_p|R\rangle. \tag{S33}$$

Since the modes and $R$ are jointly Gaussian[1], the conditional means are given by [22]

$$\langle X_p(\Delta t)|R(0){=}R\rangle = \frac{\langle X_p(\Delta t)R(0)\rangle}{\langle R^2\rangle}R. \tag{S34}$$

Using the independence of modes and Eq. (S25) and Eq. (S27), we have that

$$\langle X_p(t)R(0)\rangle = -4\langle X_p(t)X_p(0)\rangle = -4\langle X_p^2\rangle e^{-t/\tau_p}. \tag{S35}$$

[1]Note that in $d$ dimensions, the $i$th component of the vector $\mathbf{X}_p$ is jointly Gaussian with the $i$th component of the vector $\mathbf{R}$, so the equation would still hold.

Finally, by the equipartition theorem $\langle X_p^2\rangle = \frac{k_B T}{k_p} = \tilde{A}\langle R^2\rangle p^{-2}$, where $\tilde{A}$ is a constant of order unity[2]. Using this result, we find

$$\langle \Delta X_p | R\rangle = \frac{4R}{\langle R^2\rangle}\langle X_p^2\rangle (1 - e^{-\Delta t/\tau_p}) = \frac{4R\tilde{A}}{p^2}(1 - e^{-\Delta t/\tau_p}). \tag{S36}$$

To evaluate the squared terms, we need to consider terms of form $\langle X_p(t)^2 | R(0){=}R\rangle$. Since the modes and $R$ are jointly Gaussian, we have that [22]

$$\langle X_p(t)^2 | R(t'){=}R\rangle = \langle X_p^2\rangle + \frac{\langle X_p(t)R(t')\rangle^2}{\langle R^2\rangle}\left(\frac{R^2}{\langle R^2\rangle} - 1\right). \tag{S37}$$

This implies that Eq. (S30) can be expanded as

$$\langle \Delta X_p^2 | R\rangle = \left(\langle X_p^2\rangle + 16\frac{\langle X_p^2\rangle^2}{\langle R^2\rangle}\left(\frac{R^2}{\langle R^2\rangle} - 1\right)\right)(1 - 2e^{-\Delta t/\tau_p}) \tag{S38}$$

$$+ \langle X_p^2\rangle + 16\frac{(\langle X_p^2\rangle e^{-\Delta t/\tau_p})^2}{\langle R^2\rangle}\left(\frac{R^2}{\langle R^2\rangle} - 1\right), \tag{S39}$$

where we again used Eq. (S35) and Eq. (S27). Simplifying and gathering terms, we find

$$\langle \Delta X_p^2 | R\rangle = 2\langle X_p^2\rangle\left(1 - e^{-\Delta t/\tau_p}\right) + 16\frac{\langle X_p^2\rangle^2}{\langle R^2\rangle}\left(\frac{R^2}{\langle R^2\rangle} - 1\right)\left(1 - 2e^{-\Delta t/\tau_p} + e^{-2\Delta t/\tau_p}\right). \tag{S40}$$

$$= 2\frac{\tilde{A}\langle R^2\rangle}{p^2}\left(1 - e^{-\Delta t/\tau_p}\right) + 16\frac{\tilde{A}^2}{p^4}\left(R^2 - \langle R^2\rangle\right)\left(1 - e^{-\Delta t/\tau_p}\right)^2. \tag{S41}$$

To summarize, Eq. (S36) and Eq. (S41) describe the scalings of all terms of Eq. (S32). We find contributions of two types. There are terms that do not scale with $R$. All of these are absorbed into the constant $\langle \Delta R^2 | R{=}0\rangle$. The remaining terms all scale with $(4\tilde{A}R(1 - e^{-\Delta t/\tau_p})/p^2)^2$. We hence find

$$\phi_{\text{Rouse}}(R) = \langle \Delta R^2 | R\rangle - \langle \Delta R^2 | 0\rangle \sim R^2\left(\sum_{p\text{ odd}} \frac{1 - e^{-\Delta t/\tau_p}}{p^2}\right)^2. \tag{S42}$$

Finally, we split the sum into two parts; slow modes with $\Delta t/\tau_p < 1$, satisfying $p < p_c \sim \sqrt{\tau_1/\Delta t}$, and fast modes with $p > p_c$. For slow modes, we Taylor expand the exponential to leading order:

$$\sum_{p\text{ odd}<p_c} \frac{1 - e^{-\Delta t/\tau_p}}{p^2} \approx \sum_{p\text{ odd}<p_c} \frac{\Delta t/\tau_p}{p^2} \sim \frac{\Delta t}{\tau_1}\frac{p_c}{2} \sim \sqrt{\frac{\Delta t}{\tau_1}}, \tag{S43}$$

[2]This follows from noting that $k_p^{-1} \sim N\ell_0^2 \sim \langle R^2\rangle$.

where we used that $\tau_p = \tau_1/p^2$. For fast modes, by contrast, we have that

$$\sum_{p\ \mathrm{odd}>p_c}^{N} \frac{1-e^{-\Delta t/\tau_p}}{p^2} \sim \int_{p_c}^{\infty} \frac{1}{p^2} dp = \frac{1}{p_c} \sim \sqrt{\frac{\Delta t}{\tau_1}}. \tag{S44}$$

This approximation holds as long as $1 \ll p_c \ll N$. Hence, to leading order in $\Delta t$, the sum in Eq. (S42) scales with $\sqrt{\Delta t/\tau_1}$. We hence find

$$\phi_{\mathrm{Rouse}}(R) \sim R^2 \frac{\Delta t}{\tau_1}. \tag{S45}$$

Notably, the scaling with $R^2$ is expected to hold for all $\Delta t$. The temporal scaling, however, is expected to break down as higher order corrections in $\Delta t/\tau_p$ become more relevant.

Finally, we note that in three dimensions, we can decompose

$$\langle \Delta R^2 | \mathbf{R} \rangle = \langle \Delta R_x^2 | \mathbf{R} \rangle + \langle \Delta R_y^2 | \mathbf{R} \rangle + \langle \Delta R_z^2 | \mathbf{R} \rangle. \tag{S46}$$

The 1D calculation then applies to each dimension independently, after we note that the $i$:th component of $\mathbf{X}_p$ only depends on $R_i$. Summing up the contributions from each dimension then yields

$$\phi_{\mathrm{Rouse}}(\mathbf{R}) \sim |\mathbf{R}|^2 \frac{\Delta t}{\tau_1}. \tag{S47}$$

###### 4.2.2 Derivation of $\phi$ in models with spatially correlated noise

In the presence of SCFs, we expect the spatial dependence of $\langle \Delta R^2 | R \rangle$ to be dominated by the covariance of the fluctuations. We note that spatial noise correlations couple the dynamics of Rouse modes, and therefore an exact analysis as above is impossible. Instead, we construct scaling arguments for $\langle \Delta R^2 | R \rangle$ based on a phenomenological model of two particles experiencing spatially correlated noise. We first consider two Brownian particles, and then extend the analysis for fractional Brownian Motion (fBM), which describes sub-diffusive dynamics.

**Brownian motion.** First, to build intuition, we consider two Brownian particles, with $\beta = 1$, in a spatially correlated noise field. In this case, the motion of each particle is described by

$$dr_i = \sigma_1 dB_i, \tag{S48}$$

where $\sigma_1$ is the single-locus noise magnitude, and $dB_i$ is an instantaneous noise increment: $\langle dB_i(t) dB_i(t') \rangle = \delta(t, t')$. We expect that

$$\langle \Delta R^2 | R \rangle = 2\sigma_1^2 \Delta t - 2\sigma_1^2 \int_0^{\Delta t} \int_0^{\Delta t} \langle dB_1(t) dB_2(t') | R \rangle dt\, dt'. \tag{S49}$$

We assume that noise correlations are also instantaneous ($\langle dB_1(t) dB_2(t') | R \rangle \sim \delta(t, t')$) and that they decay spatially according to $C_{\mathrm{cor}}(R)$ without an explicit time-dependence.

If the time-interval is sufficiently short that the correlation magnitude remains nearly constant (with exponentially decaying correlations, this is the case if $\Delta R \ll \lambda$), we then expect

$$\phi_{\mathrm{SCF}}(R) = \langle \Delta R^2 | R \rangle - \langle \Delta R^2 | 0 \rangle \sim (C_{\mathrm{cor}}(0) - C_{\mathrm{cor}}(R))\Delta t. \tag{S50}$$

This is consistent with $\phi \sim \Delta t^{\kappa}$ for $\kappa = \beta = 1$ in the case of diffusive motion.

**Fractional Brownian Motion.** Chromosomal loci display sub-diffusive motion; hence, their dynamics cannot be described as simple Brownian motion. In the case of uncorrelated noise, we could describe $R$ as a fractional fBM process with Hurst parameter $H = 1/4$ [23, 24], expressed as a Weyl integral

$$R(t) = R(0) + \int_0^t K_H(t,s)\sigma dB(s) \tag{S51}$$

where $dB(s)$ is a Brownian noise increment, and $K_H$ is a kernel defined by polymer dynamics. Intuitively, the kernel couples the current dynamics of $R$ to noise at a time $t - s$ in the past. $dB(s)$ should be seen as a linear combination of noise terms experienced by all monomers at time $s$, weighted appropriately. With $t - s > 0$ fixed, in the limit of $t, s \to \infty$, we have

$$K_H(t,s) \sim (t-s)^{H-1/2}. \tag{S52}$$

This kernel gives rise to the required scaling of $\langle (R(t) - R(s))^2 \rangle \sim (t-s)^{2H}$ [23].

We now suppose that the addition of SCFs does not change the MSD scalings of the distance $R$, and that the kernel displays the same scalings as above. We then assume that the noise amplitudes depend on the current value of $R(s)$. We write $R$ as an Itô integral:

$$R(t) = R(0) + \int_0^t K_H(t,s)\sigma(R(s))dB(s). \tag{S53}$$

Here, similar to the Brownian case, we assume that $\sigma^2 = 2\sigma_1^2 - 2C_{\mathrm{cor}}(R)$, where $\sigma_1$ is the standard deviation of single-locus increments, and $C_{\mathrm{cor}}(R)$ is a smooth function that describes the decay of spatial noise correlations. Since $\lim_{R\to\pm\infty} C_{\mathrm{cor}}(R) = 0$, $\sigma(R)$ is bounded. We are interested in the leading order behavior of $\langle \Delta R^2 | R \rangle$ as $\Delta t \to 0$.

First, we write

$$\Delta R = \int_t^{t+\Delta t} K_H(t+\Delta t, s)\sigma(R(s))dB(s) + \int_0^t [K_H(t+\Delta t, s) - K_H(t,s)]\sigma(R(s))dB(s) \tag{S54}$$

Since the integrals are non-overlapping, and since $K_H$ can only depend on the previous states of the system, by the Itô isometry

$$\langle \Delta R^2 | R \rangle = \underbrace{\int_t^{t+\Delta t} K_H(t+\Delta t, s)^2 \langle \sigma(R(s))^2 | R \rangle ds}_{I_1} \tag{S55}$$

$$+ \underbrace{\int_0^t [K_H(t+\Delta t, s) - K_H(t,s)]^2 \langle \sigma(R(s))^2 | R \rangle ds}_{I_2}. \tag{S56}$$

To estimate the scaling behavior of $I_1$, we assume that $t$ is sufficiently large for Eq. (S52) to hold. We also again assume that $\Delta t$ is sufficiently small that in the interval $[t, t + \Delta t]$ $\langle \sigma(R(s))^2 | R \rangle \approx \sigma(R)^2$. We then find

$$I_1 \sim \sigma(R)^2 \int_t^{t+\Delta t} (t + \Delta t - s)^{2H-1} ds = \sigma(R)^2 \frac{\Delta t^{2H}}{2H}. \tag{S57}$$

Notably, the term follows the same $\Delta t^{2H}$ scaling as the MSD of $R$.

Next, we estimate $I_2$. We fix $\Delta t < \epsilon \ll t$ and split the integral in two parts:

$$I_2 = \int_0^{t-\epsilon} [K_H(t + \Delta t, s) - K_H(t, s)]^2 \langle \sigma(R(s))^2 | R \rangle ds \tag{S58}$$

$$+ \int_{t-\epsilon}^{t} [K_H(t + \Delta t, s) - K_H(t, s)]^2 \langle \sigma(R(s))^2 | R \rangle ds. \tag{S59}$$

In the first interval $[0, t - \epsilon]$ the kernel is differentiable, so that we can Taylor expand $K_H(t + \Delta t, s)$. In the second interval, we again use the large $t, s$ scaling limit of the kernel, and approximate $\langle \sigma(R(s))^2 | R \rangle \approx \sigma(R)^2$.

$$I_2 \approx \int_0^{t-\epsilon} \left[ \frac{\partial K_H(t, s)}{\partial t} \Delta t \right]^2 \langle \sigma(R(s))^2 | R \rangle ds \tag{S60}$$

$$+ A\sigma(R)^2 \int_{t-\epsilon}^{t} [(t + \Delta t - s)^{H-1/2} - (t - s)^{H-1/2}]^2 ds, \tag{S61}$$

where $A$ is a constant of order unity. For fixed $\epsilon$, the first term is order $\Delta t^2 \ll \Delta t^{2H}$, and hence negligible compared to $I_1$. By comparing the second term to $I_1$, we expect it to give a smaller correction with the same scaling. To see this, we change variable to $\tilde{s} = (t - s)/\Delta t$:

$$I_2 \approx A\sigma(R)^2 \Delta t^{2H} \int_0^{\epsilon/\Delta t} [(\tilde{s} + 1)^{H-1/2} - \tilde{s}^{H-1/2}]^2 d\tilde{s} + \mathcal{O}(\Delta t^2). \tag{S62}$$

For fixed $\epsilon$, as $\Delta t \to 0$, the integral converges to a constant.

By combining Eq. (S57) and Eq. (S62), we hence find that in the limit of $t \gg 0$, $\Delta t \to 0$, to leading order

$$\langle \Delta R^2 | R \rangle \sim \sigma(R)^2 \Delta t^{2H} = 2(\sigma_1^2 - C_{\text{cor}}(R)) \Delta t^{2H}. \tag{S63}$$

Since $2H = \beta$ sets the dynamic exponent, we hence find

$$\phi_{\text{SCF}}(R) = \langle \Delta R^2 | R \rangle - \langle \Delta R^2 | 0 \rangle \sim (C_{\text{cor}}(0) - C_{\text{cor}}(R)) \Delta t^{\beta}, \tag{S64}$$

as was to be demonstrated. Our result relies on the following approximations:

1. We assumed that $t$ is sufficiently large, so that the finite history of the process does not affect the kernel. This should hold if the system has had time to reach a steady-state.

2. We assumed that $\Delta t$ is sufficiently short that the correlation magnitude remains approximately constant.

3. We assumed that the kernel $K_H(t, s)$ is set by polymer dynamics. This seems reasonable for experimental data, where the mean-squared distances consistently follow a $t^{1/2}$ scaling.

4. We assumed that the noise increments appear uncorrelated on the relevant time-scales. If the noise was also temporally correlated on time scales comparable to the relaxation time of the polymer segment, we might expect a different scaling with $\Delta t$. Temporal noise correlations would also cause the correlation magnitude to vary slower in time, so that Approximation 2 would hold for larger $\Delta t$.

##### 4.3 Transition from SCF to Rouse scaling

The different $\Delta t$-scalings of the Rouse and SCF contributions ($\Delta t$ vs. $\Delta t^{1/2}$) imply that, in the presence of both effects, $\phi \sim \phi_{\text{SCF}}$ for small $\Delta t$, whereas for large $\Delta t$, we expect a cross-over to the Rouse-scaling. Here, we provide a simple scaling argument for the minimum time-scale $\Delta t$ required for the $\phi(R; \Delta t)$ curves to show a plateau in the presence of SCFs. This scaling relation can be used to check whether an observed Rouse-like $\phi$ scaling could be due to the imaging interval $\Delta t$ being too long for a given genomic length $s$.

For a polymer with spatially correlated noise, we expect the SCF and Rouse contributions to $\phi$ to be of similar magnitudes when

$$R^2 \Delta t / \tau_s \sim C_0 (1 - e^{-R/\lambda}) \Delta t^{\beta}. \tag{S65}$$

This follows from equating Eq. (S45) and Eq. (S64) We note that $C_0 \Delta t^{\beta} \leq 2\langle \Delta r \rangle$, where equality would imply that at $R = 0$, the two loci would move in a perfectly correlated way, and at $R \gg \lambda$, they would move independently. Hence, up to a constant of order unity, we expect a cross-over from SCF- to Rouse-scalings at a crossover distance $R_c$ when

$$\Delta t / \tau_s \sim 2\langle \Delta r \rangle (1 - e^{-R_c/\lambda}) R_c^{-2}. \tag{S66}$$

For $\tau_s \sim s^2$, we hence see that for a larger $s$, a larger $\Delta t$ can be used. Alternatively, for fixed $\Delta t$, the crossover distance $R_c$ grows with $s$.

Next, we ask under what conditions the plateau in $\phi$, characteristic of SCFs, can be resolved. This requires that $\Delta t$ is sufficiently small that the cross-over happens at distances $R_c > \lambda$. The exact threshold $R_{\text{thresh}}$ will depend on the statistical accuracy of the experiment; with fewer samples, a larger threshold should be chosen, whereas with high statistical accuracy, a lower threshold may be used. In either case, it should be noted that the $\phi(R)$ curve will start to deviate from the SCF scaling for $R > R_c$.

Based on the experimental data for fly embryos (SI Fig. S12, S7), $\lambda \approx 0.4$ µm, and the plateau is clearly visible around 2 µm. Hence, resolving the plateau given $\Delta t = 28$ s and the experimentally measured $\langle \Delta r^2 \rangle \approx 0.3$ µm$^2$ would require that

$$\tau_s > \Delta t \frac{R_{\text{tresh}}^2}{2\langle \Delta r^2 \rangle} \approx 186 \text{ s}. \tag{S67}$$

Based on previous calculations [2], this is true for the experimental data sets with $s >$ 100 kb. Hence, we only show data for $s >$ 100 kb in Main Fig. 2g. For shorter genomic separations, we confirm that fluctuation-distance profiles still appear concave, but the data do not extend far enough in $R$ to clearly plateau, and hence to reliably fit the data (Fig. S7a). SCF simulations show similar results (Fig. S7b). By contrast, in Rouse simulations, even for $s <$ 100 kb, we observe $\phi \sim R^2$ scalings (Fig. S7c).

To summarize, the different scalings of $\phi_{\text{Rouse}} \sim \Delta t$ and $\phi_{\text{SCF}} \sim \Delta t^{1/2}$ imply that for large $\Delta t$, Rouse-like effects will start to dominate $\phi$, even if SCFs are present. For given $\Delta t$ and $\tau_s$, one can calculate the distance at which Rouse-like scaling will start to dominate. If this distance is sufficiently large compared to the correlation length $\lambda$, fluctuation-distance profiles can be used to accurately test for SCFs, as well as to quantify the level at which $\phi$ plateaus. If the cross-over distance is smaller than or close to $\lambda$, one may still be able to observe convex behavior of the fluctuation-distance profile at small $R$, but at large $R$ the scaling will switch over to a Rouse-like $\phi \sim R^2$ scaling.

#### 4.4 Collapse of fluctuation-distance profiles for the Rouse model

Our theory predicts that for $\Delta t \ll \tau_s$, we expect fluctuation-distance profiles to collapse for $\phi/\Delta t \sim (Rs^{-\gamma/2})^2$, where $\gamma$ defines the scaling $\tau_s \sim s^\gamma$. We confirm that at short time-scales $\Delta t = 1 \ll \min(\tau_s) \approx 100$ s [2], the Rouse model with or without confinement shows the expected scalings for $\gamma = 2$ (Fig. S8b).

At $\Delta t \approx 28$ s intervals used in experiments, the data somewhat deviate from the expected scaling. We still observe a $\phi \sim R^2$ scaling and a clear dependence on $s$ (Fig. S8a), unlike in SCF simulations. However, the relatively large $\Delta t$ value means that our expansion in $\Delta t/\tau_s$ (SI Appendix 4.2) breaks down.

Finally, to test the $\phi \sim \Delta t$ scaling, we plot $\phi \Delta t^{-\kappa}$ for varying $\Delta t$. For experimental data, the best collapse is achieved with $\kappa = 1/2$, as expected in the presence of SCFs (Fig. S10a,b). By contrast, for the Rouse model, the data collapse for $\kappa = 1$, as predicted (Fig. S10d).

#### 4.5 Spatially correlated equilibrium white noise does not recapitulate experimental data

As mentioned in the main text, we tested whether an equilibrium model with spatially correlated white noise could explain the experimental observations.

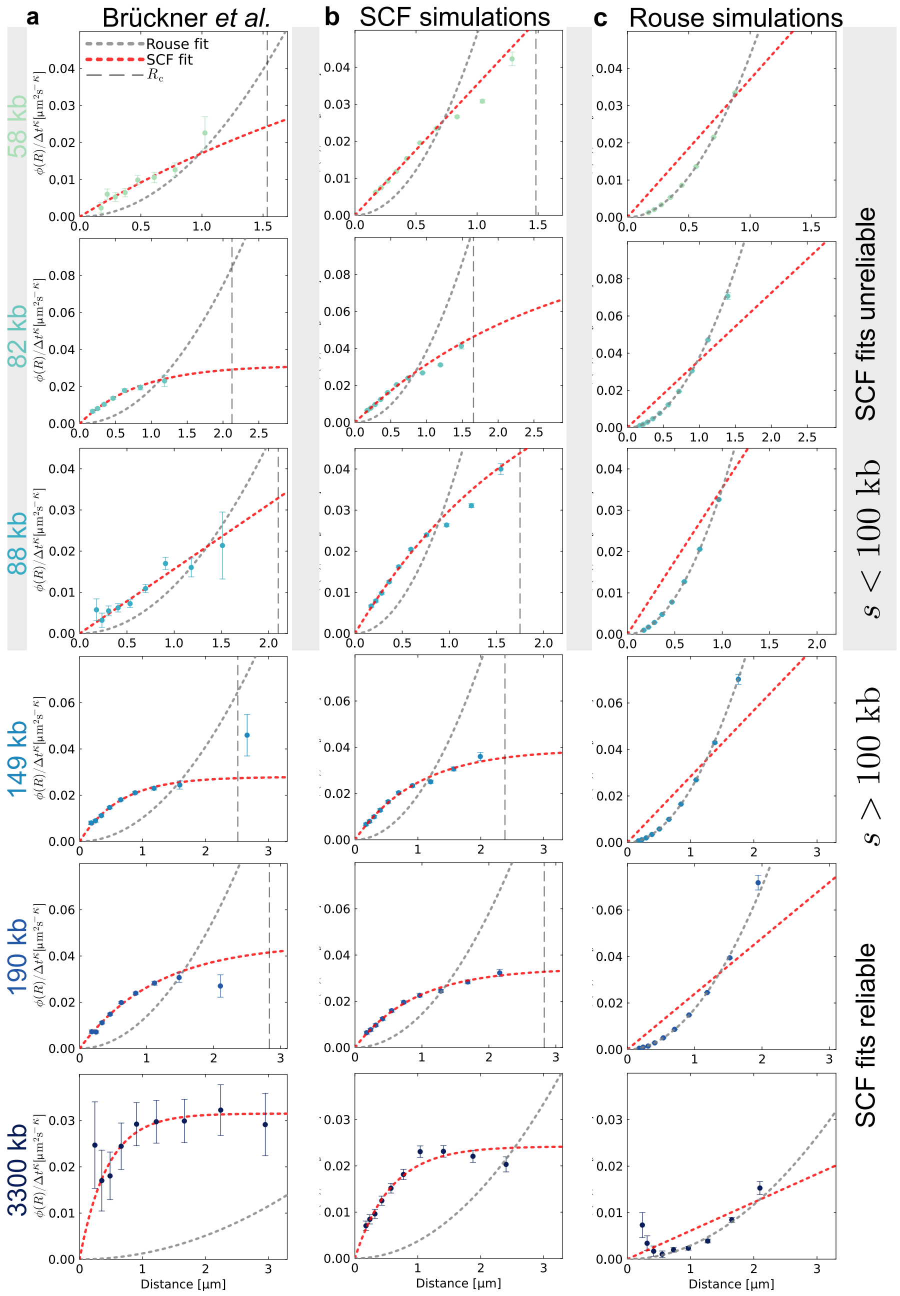

Figure S7: **Individual fluctuation-distance profiles.** $\phi(R)$ curve estimates for... **(a)** fruit fly embryo data [2]. Vertical dashed lines indicate distances at which we estimate Rouse-like scalings to start to dominate. For $s < 100$ kb, the data show concave behavior, but do not extend far enough in $R$ to be reliably fitted with the SCF prediction. For $s > 100$ kb, the plateau can be seen more clearly. In all cases, the SCF prediction (red line) has a smaller error than the Rouse prediction (gray line) when fitted to the data. Note that the individual curves for $s = 595$ kb are shown in Main Fig. 2. **(b)** SCF simulations. Similar to experiments, the curves are concave, but do not always extend far enough in $R$ to show a clear plateau. **(c)** Rouse simulations. For all $s$, the data now appear convex, and can be fitted with the Rouse-prediction for $\phi \sim R^2$.

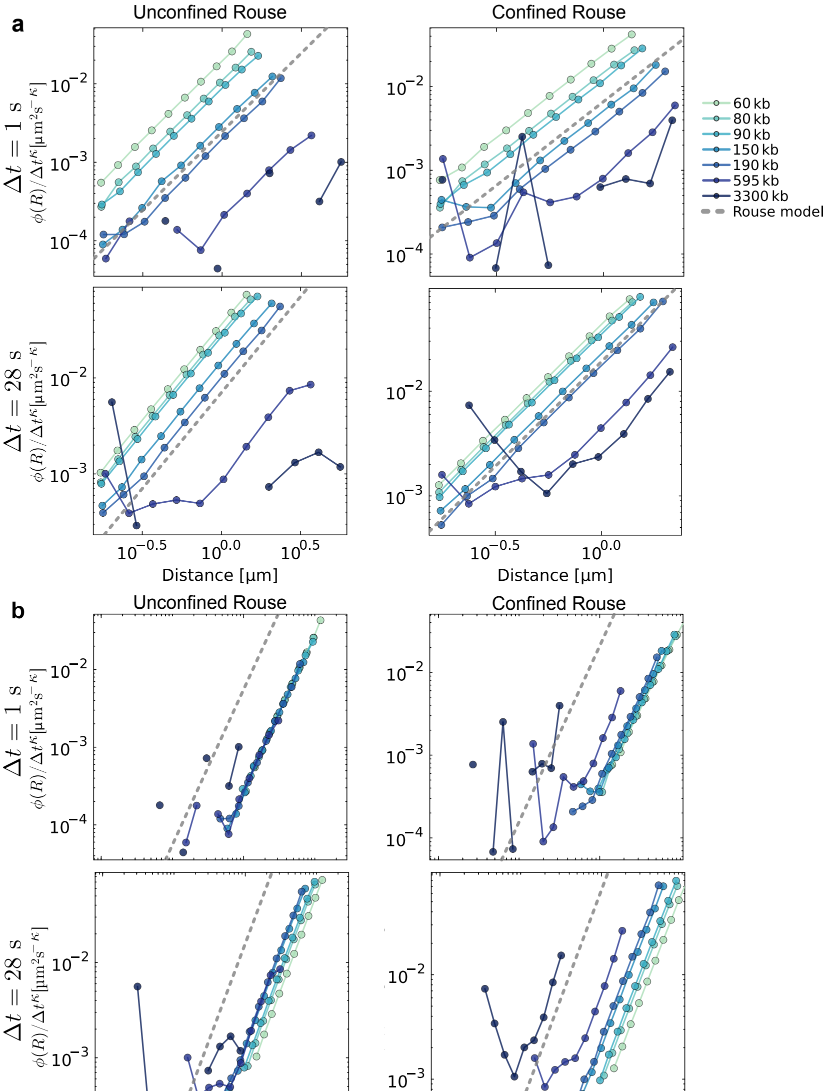

Figure S8: **Fluctuation-distance profile collapses for Rouse simulations.** **(a)** Table of log-log plots of $\phi(R)$ vs $R$ curves for the unconfined and confined Rouse models, at $\Delta t = 1$ or 28 s. The data show a clear $\phi \sim R^2$ scaling, as indicated by the gray dashed line. **(b)** Similar to (a), but x-axis scaled by $s^{-1}$. The data show a clear collapse at short time-scales $\Delta t = 1$ s, as expected based on the theoretical prediction $\phi_{\text{Rouse}} \sim R^2 s^{-\gamma/2}$, with $\gamma = 2$ for the Rouse model. At longer time-scales, the data somewhat deviate from the collapse.

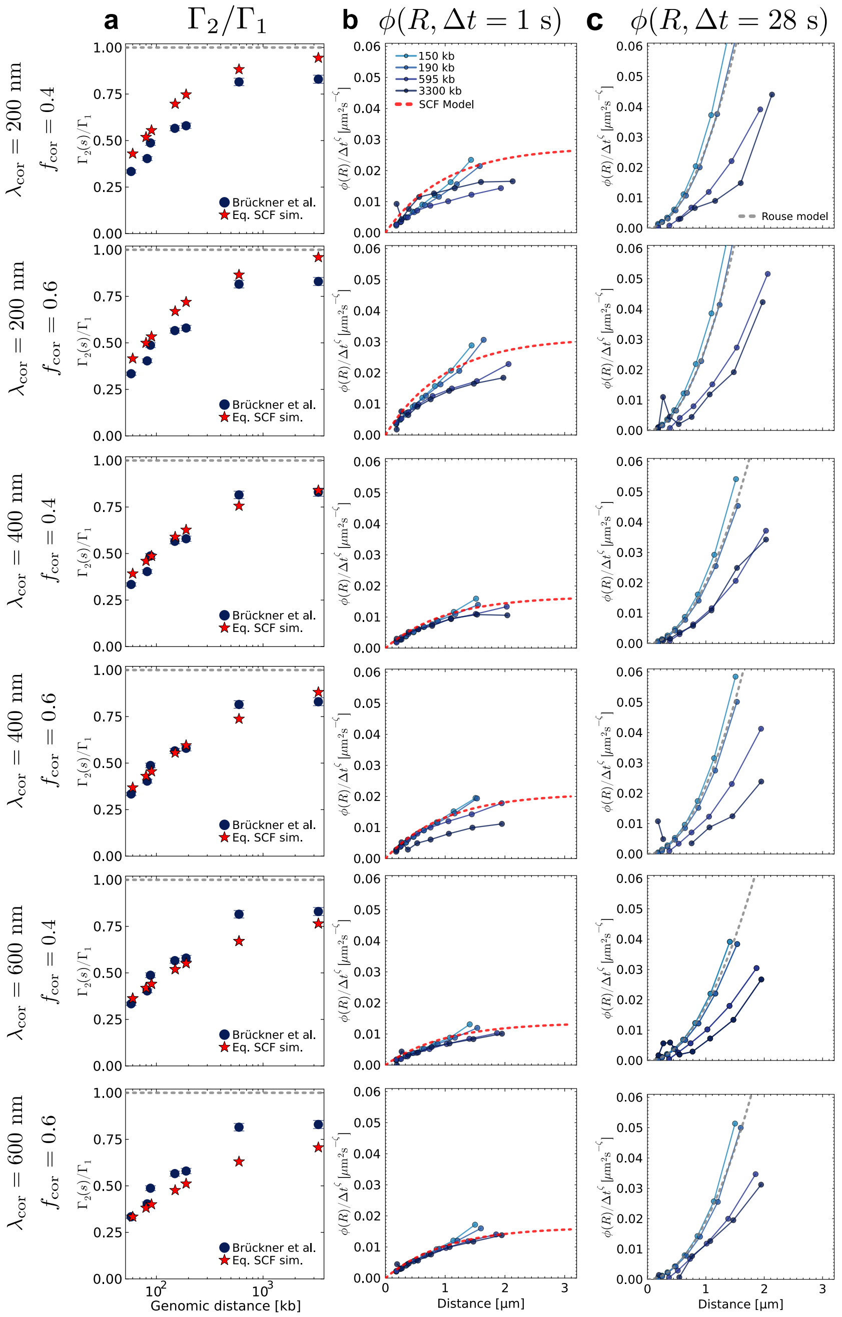

Figure S9: **(a)** $\Gamma_2/\Gamma_1$ curves for simulations with spatially correlated equilibrium white noise, for varying $f_{\text{cor}}$ and $\lambda_{\text{cor}}$. **(b)** $\phi(R)$ curves for the same models, calculated with $\Delta t \approx 1$ s. At these short time-scales most models show the expected scaling of $\phi \sim 1 - e^{-R/\lambda}$ expected for SCFs. **(c)** $\phi(R)$ curves calculated at the temporal resolution of $\Delta t \approx 28$ s used in experiments [2]. At these longer time-scales, the equilibrium simulations do not show the experimentally observed SCF scaling.

The free parameters of the model are $f_{\text{cor}}$, the fraction of correlated noise, and $\lambda_{\text{cor}}$, the length scale across with noise is correlated. For a range of values, we find that the equilibrium SCF model is able to recapitulate the experimentally observed $\Gamma_2(s)$ curves (Fig. S9a).

Next, we calculate $\phi$ for the model with equilibrium SCFs. We find that at short time-scales ($\Delta t = 1$ s), the $\phi(R, \Delta t)$-curves display a plateau independent of $s$, as expected based on theory (Fig. S9b). However, at the experimentally sampled time-scales ($\Delta t \approx 28$ s), the scaling follows the Rouse-like $\phi \sim (R/s)^2$ form (Fig. S9c). Adjusting $\lambda_{\text{cor}}$ or $f_{\text{cor}}$ while keeping $\Gamma_1$ constant does not result in a better match between experiments and simulations. Hence, although the model follows our theoretical predictions as $\Delta t \to 0$, it fails to predict the shape of the fluctuation-distance profiles at experimentally probed time-scales. We hence conclude that the equilibrium white noise correlations do not explain the experimental data.

Why does the model's $\phi$-scaling break down at $\Delta t = \mathcal{O}(10\text{ s})$? The values for $\Gamma_2$ vary between 0.022 and 0.044 $\mu\text{ms}^{-1/2}$ for $s = 58$ kb and $s = 3300$ kb correspondingly. This means that the spatial distances between the loci are expected to change by 240-470 nm in an interval $\Delta t = 28$ s. This length scale is comparable to the estimated correlation length $\lambda_{\text{cor}} \approx 500$ nm, which is largely set by the scalings of $R(s)$ and $\Gamma_2(s)$ (Fig. S9). Hence, in the interval $\Delta t \approx 28$ s, the distance between loci changes so much that SCF effects can appear averaged out, breaking assumption 2 introduced in Section 4.2.2. In non-equilibrium flow simulations, this effect is mitigated by the temporal correlations of the flow field.

### 5 Data processing

#### 5.1 Calculating MSDs, mean-squared distances, and diffusion coefficients

For all experimental data sets, we calculated the single and two-locus mean-squared displacement curves as the autocorrelation curves of the 2D change in position or inter-locus distance. $z$-displacements were not used for experimental data due to the larger experimental error in this dimension. The curves were rescaled by 3/2, assuming isotropy across dimensions. For consistency with the large $t$-limit of the two-locus MSD curves, the mean-squared distance between loci was also calculated in 2D, and then rescaled by 3/2. Our analysis could also be performed using 3D data, but the relative error of $\Gamma_2$ would be larger.

Given the MSD curves, we calculated $\Gamma_1 = \langle \Delta r^2 \rangle \Delta t^{-1/2}$ from the MSD between consecutive frames [19]. The error estimate for $\Gamma_1$ was calculated from the standard error of the mean.

For $\Gamma_2(s)$, we calculated $\Gamma_2(s) = 1/2 \langle \Delta R^2 \rangle_s \Delta t^{-1/2}$ from the two-locus MSD between consecutive frames. The error estimate was again calculated from the standard error of the mean. With these definitions, if locus motion were uncorrelated, we would find $\Gamma_2 = \Gamma_1$.

For ratios $\Gamma_2/\Gamma_1$ (Main Fig. 1d), we propagated the errors by adding the relative errors of $\Gamma_2$ and $\Gamma_1$.

##### 5.2 Effect of subdiffusion exponent on results

To calculate $\Gamma_2$ and $\Gamma_1$ as described in Appendix 5.1, the subdiffusion exponent $\beta$ needs to be determined. By fitting the MSD, we find $\beta \approx 1/2$ (Main Fig. 1b) [2]. However, experiments suggest that the subdiffusive exponent might vary across species and timescales [1]. We now explain why our results do not qualitatively depend on $\beta$.

Using a different value $\beta'$ would only change our $\Gamma$ estimates by a factor of $\Delta t^{-\beta'+1/2}$. Hence, when comparing results from experiments with the same $\Delta t$, such as in Main Fig. 1, using a different value of $\beta$ would not qualitatively change our findings.

When comparing $\Gamma_2/\Gamma_1$ curves, such as in Main Fig. 1d, the factors of $\Delta t^{-\beta}$ cancel. Therefore, these results are unaffected by the choice of $\beta$.

Finally, for plots of $\phi(R, \Delta t)$ or $\langle \Delta R^2 | R \rangle$, we make no assumptions about $\beta$. Our analysis shows (SI Appendix 4.2) that for the Rouse model, corrections to $\phi(R)$-curves scale linearly with $\Delta t$ (Eq. 2). By contrast, with SCFs, we expect corrections to scale with $\Delta t^{\beta}$. For experimental data from [2, 4], we find that data collapse for $\phi \sim \Delta t^{\kappa}$, with $\kappa = 1/2$, consistent with $\beta = 1/2$ (Fig. S10a,b).

##### 5.3 Estimating $\phi(R)$ from data

To calculate $\phi(R)$, we require estimates of the conditional average $\langle \Delta R^2 | R \rangle$ as a function of distance, and the extrapolated value $\langle \Delta R^2 | R{=}0 \rangle$. We proceed as follows:

- **Binning.** For each genomic separation $s$, we calculate pairs $\{R, \Delta R_\alpha^2\}$, where $\alpha$ is the spatial dimension. For experimental data, we discard points with $R < \sigma_{\text{exp}}$ (Table S6) and use only $\alpha \in \{x, y\}$, since $z$-localization errors are larger. We logarithmically bin $\Delta R_\alpha^2$ by $R$, merging edge bins with fewer than 30 samples. Results are robust to the choice of binning method and bin number (Fig. S11). Plots of $2\langle \Delta r^2 \rangle - \langle \Delta R^2 | R \rangle$ suggest that fluctuation magnitudes decay roughly exponentially (Fig. S12b).
- **Fitting.** We fit the binned data using weighted least squares (Julia LsqFit), with weights given by the inverse squared standard error of the mean. We compare two models: an exponential saturation consistent with SCFs,

$$\langle \Delta R^2 | R \rangle = A + 2C(1 - e^{-R/\lambda}), \tag{S68}$$

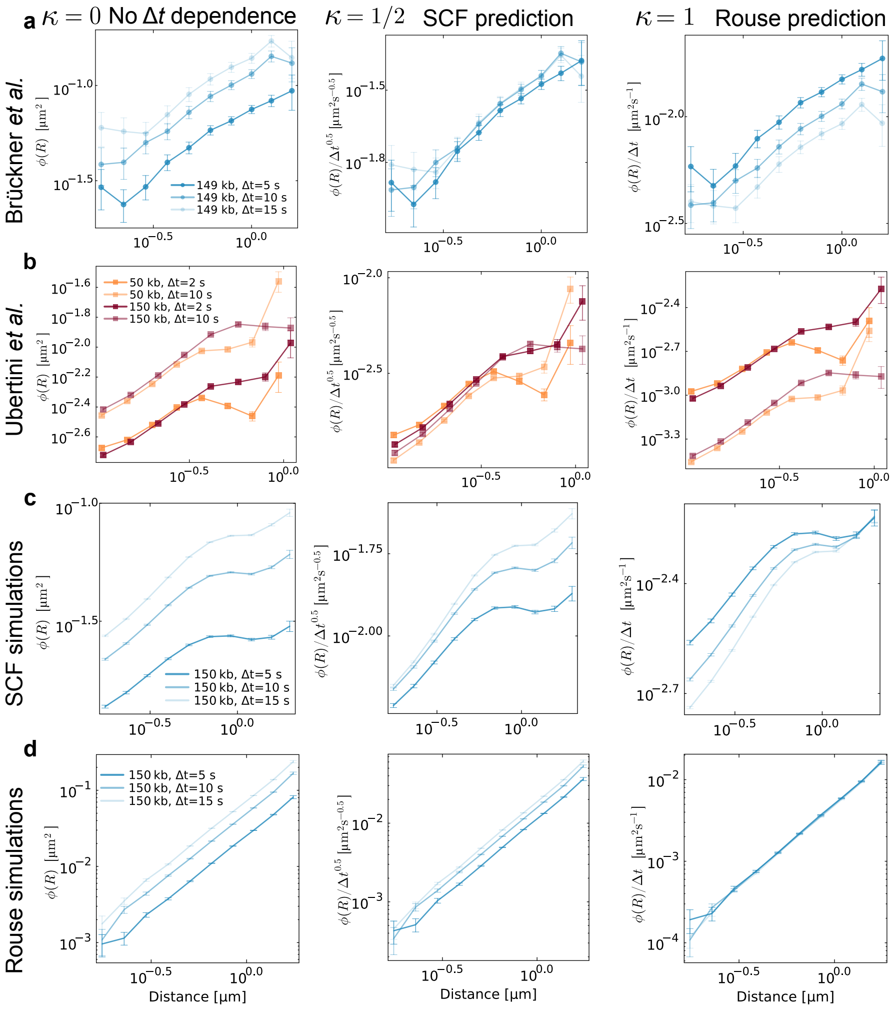

Figure S10: **Testing $\phi\Delta t^{-\kappa}$ collapses.** A table of $\phi\Delta t^{-\kappa}$ plots for $\kappa \in \{0, 1/2, 1\}$. $\kappa = 0$ would correspond to $\phi$ being independent of $\Delta t$. $\kappa = \beta \approx 1/2$ is the prediction for SCFs. $\kappa = 1$ is the prediction for the Rouse model. **(a)** Row shows fruit fly embryo data from [2]. The data show the best collapse with $\kappa = 1/2$, as expected in the presence of SCFs. **(b)** mESC data from [4]. Again, $\kappa = 1/2$ gives the best collapse. **(c)** Data from SCF simulations. At low $R$ and $\Delta t$, $\kappa = 1/2$ gives the best collapse. For large $R$, where Rouse-scalings are expected to dominate (SI Appendix 4.3), $\kappa = 1$ gives a better collapse. **(d)** Data from Rouse simulations. As expected, $\kappa = 1$ gives the best collapse.

and a quadratic form consistent with the Rouse model,

$$\langle \Delta R^2 | R \rangle = A' + BR^2, \tag{S69}$$

with all fit parameters constrained to be non-negative. We select the better model based on weighted root mean squared errors across varying bin numbers. For almost all experimental data sets, the exponential fit is preferred, with exceptions at $s < 100$ kb where intermediate scaling is expected (SI Appendix 4.3).

- **Extrapolation to $R = 0$.** We estimate $\langle \Delta R^2 | R{=}0 \rangle$ as the selected fit function evaluated at $R = 0$: $A$ for the exponential model, $A'$ for the quadratic model. Errors on $\phi$ are calculated as the sum of each bin's standard error and the estimated error in the $R{=}0$ extrapolation.

- **Fitting requirements.** The exponential fit requires data spanning a sufficient range in $R$ relative to the correlation length $\lambda$. When most data fall in the regime $R < \lambda$, Eq. (S68) reduces to a linear relation,

  $$\langle \Delta R^2 | R \rangle \approx (A - 2C) + 2CR/\lambda, \tag{S70}$$

  which has only two free parameters and cannot constrain the full exponential. In mESC data [3], the SCF plateau is resolved only when $R \gtrsim 1.5$ µm (Fig. S13a). Perturbations that reduce looping frequency (CTCF deletion, cohesin degradation) increase the explored range in $R$ and allow the plateau to be resolved, while conditions with frequent looping (WT, WAPL degradation, CTCF degron) restrict the $R$-range and prevent reliable fitting (Fig. S13b).

###### 5.3.1 Binning raw data

##### 5.4 Calculating encounter frequencies and durations

We use simulated time-trajectories for the locus distance $R(t)$ to calculate estimates for both the encounter frequencies and encounter durations. An encounter is defined as a distance of $R \leq 300 \times \sqrt{d/3}$ nm, motivated by the observation that transcriptional activity increases at this distance [2]. The factor $\sqrt{d/3}$, where $d$ is the dimension of the simulations (2 or 3) is chosen because for fixed spring stiffness, distances scale with $\sqrt{d}$ in the Rouse model.

At the end of each trajectory, we either have a period of time in which no encounter happens, or an encounter that lasts until the end of the trajectory. Neglecting these periods would bias sampling towards lower first passage times and encounter durations. Hence, similar to previous works [3], we use the Kaplan-Meier estimator [26] for the survival function of both the no-encounter and encounter states. The survival function $S_i(t)$ of a state $i$ is defined as the probability that the system remains in the same state for a time $\geq t$.

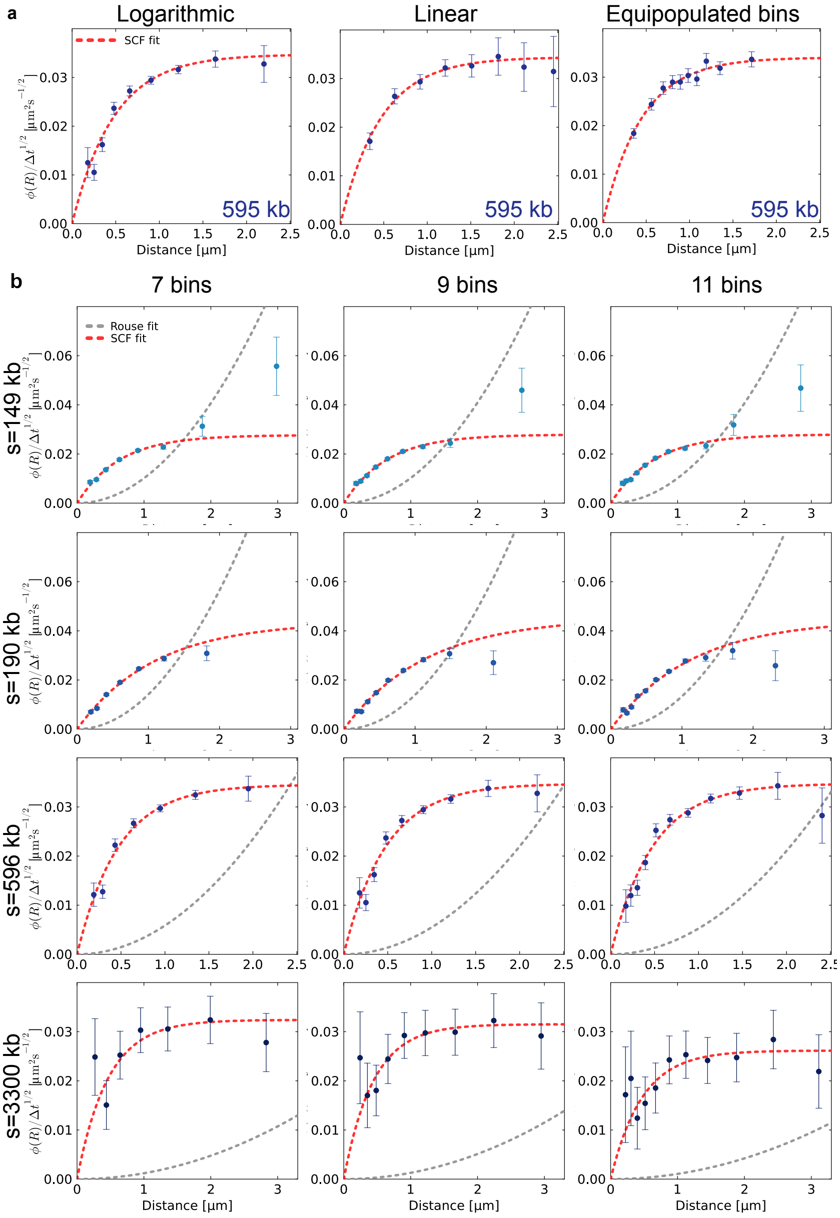

Figure S11: **Robustness of results with varying binning methods.** **(a)** Fluctuation-distance profiles calculated using logarithmic, linear, or equipopulated bins. Results are similar, and clearly show the SCF predicted plateau. **(b)** Fluctuation-distance profiles calculated using varying numbers of bins.

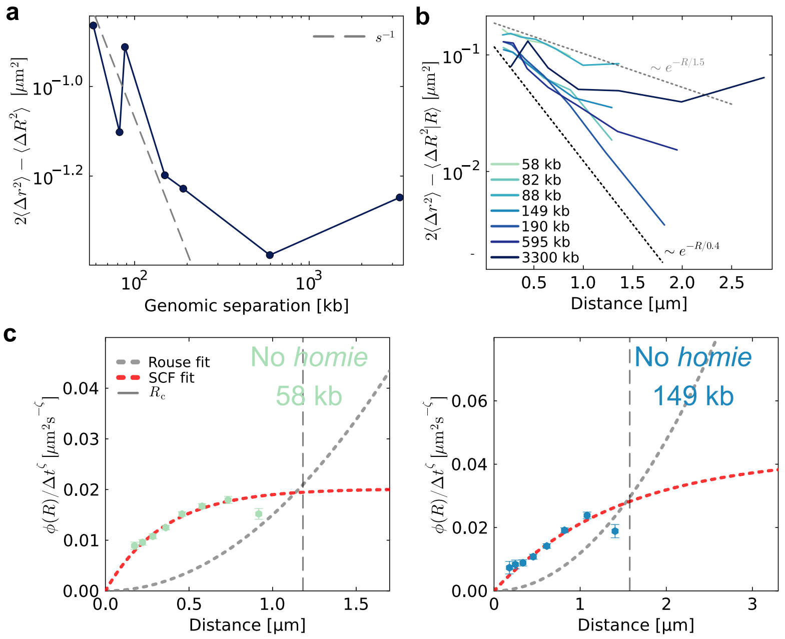

Figure S12: **Experimental scalings and no *homie* data.** **(a)** Plots of $2\langle \Delta r^2 \rangle - \langle \Delta R^2 \rangle$ against genomic separation. The differences do not follow an $s^{-1}$ scaling, as would be expected if only center of mass correlations affected the relative motion between loci [25]. **(b)** Plots of $2\langle \Delta r^2 \rangle - \langle \Delta R^2 | R \rangle$ as a function of spatial distance. Dashed lines indicate exponentially decaying trends with distance. We estimate that there is a decaying trend with distance, with an effective decay length $0.4 < \lambda < 1.5$ μm. **(c)** $\phi(R)$ curves for data without *homie* insulator pairs at the tracked loci. The data are better fitted by the SCF prediction $\phi(R) \sim 1 - e^{-R/\lambda}$ than the Rouse prediction $\phi \sim R^2$. Vertical dashed lines denotes $R_c$, the length scale at which Rouse-like scaling behavior is expected to start to dominate (SI Appendix 4.3).

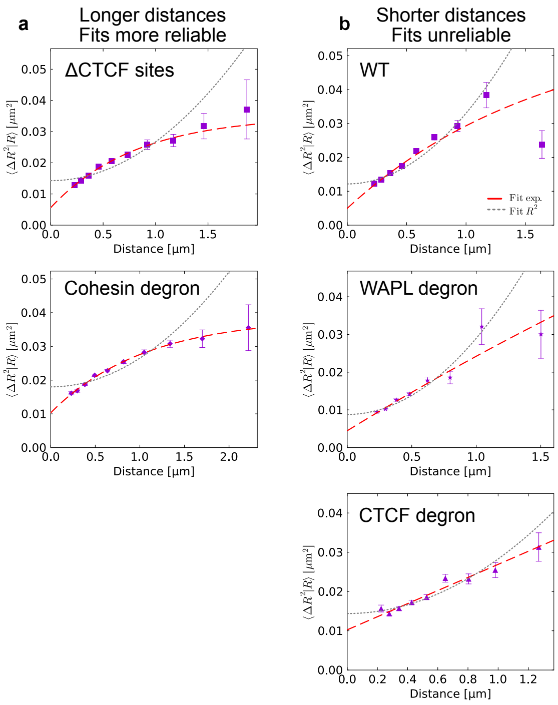

Figure S13: **Fitting Gabriele *et al.* data. (a)** $\langle \Delta R^2 | R \rangle$ curves for mESC data from [3]. Deletion of CTCF binding sites near the labeled loci, as well as cohesin degradation, increase the range of distances explored by the locus pair, because both perturbations reduce the frequency of looping. The increased range in $R$ allows the resolution of the plateau associated with SCFs. **(b)** The presence of CTCF sites (WT), the degradation of WAPL, and the CTCF degron all show a smaller range of distances. Because of this, the $\langle \Delta R^2 | R \rangle$ data cannot be reliably fitted with the SCF prediction.

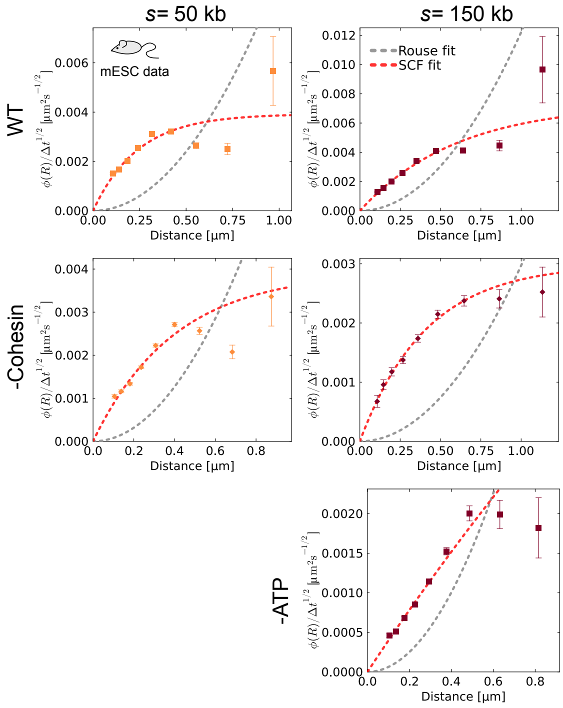

Figure S14: **Individual fluctuation-distance profiles for mESC data.** Table of curves shown in Main Fig. 3, including error bars based on the s.e.m. for each distance bin.

The median encounter duration $\tau_{\text{enc}}$ is found as the first time at which half of encounters have ended: $S_{\text{enc}}(\tau_{\text{enc}}) \geq 0.5$.

For encounter frequencies, we calculate the survival curves for the no encounter state, conditioned on the initial distance $R_0$ at $t = 0$. In practice, for each trajectory, we find all time-points at which $R = R_0$, and calculate the times until the next encounters (or the end of the trajectory). We then use these times to calculate the Kaplan-Meier estimate for the survival function.

Given the survival function for a given starting distance, the median FPT $\tau_{\text{FPT}}(R_0)$ can be found as the first time $\tau$ for which $S_{\text{no enc}}(\tau_{\text{FPT}}|R_0) \geq 0.5$. The encounter frequency is then defined as $f_{\text{enc}}(R_0) = 1/\tau_{\text{enc}}(R_0)$. In Main Fig. 4, we consider the encounter frequency from the mean distance $R_0 = \langle R \rangle$.

### 6 Supplementary Tables

| Data set | Condition | $s$ [kb] | $\lambda$ [nm] | $C$ [μm$^2$] | $C\Delta t^{-1/2}$ [μm$^2$s$^{-1/2}$] | Figure |
|---|---|---|---|---|---|---|
| Brückner *et al.* | No *homie* | 149 | $1300 \pm 470$ | $0.1 \pm 0.019$ | $0.019 \pm 0.0037$ | S12c |
| Brückner *et al.* | *homie* | 149 | $650 \pm 73$ | $0.074 \pm 0.0022$ | $0.014 \pm 0.00041$ | S7a |
| Brückner *et al.* | *homie* | 190 | $1100 \pm 240$ | $0.12 \pm 0.013$ | $0.022 \pm 0.0024$ | S7a |
| Brückner *et al.* | *homie* | 595 | $500 \pm 100$ | $0.093 \pm 0.0078$ | $0.017 \pm 0.0015$ | Main Fig. 2c |
| Brückner *et al.* | *homie* | 3300 | $460 \pm 160$ | $0.086 \pm 0.025$ | $0.016 \pm 0.0047$ | S7a |

Table S4: **Fitted values for data from fruit fly embryos.** Values were found by fitting $\langle \Delta R^2 | R \rangle$ data with the SCF prediction $\langle \Delta R^2 | R \rangle = \langle \Delta R^2 | 0 \rangle + 2C(1 - e^{-R/\lambda})$. Our theory predicts that $C \sim \Delta t^{1/2}$, so we also report scaled values. We report values for $s > 100$ kb, where the SCF scaling is expected to dominate over polymeric effects (SI Appendix 4.3).

| Data set | Condition | $s$ [kb] | $\lambda$ [nm] | $C$ [μm$^2$] | $C\Delta t^{-1/2}$ [μm$^2$s$^{-1/2}$] | Figure |
|---|---|---|---|---|---|---|
| Ubertini *et al.* | WT | 150 | $590 \pm 64$ | $0.0051 \pm 0.00033$ | $0.0036 \pm 0.00024$ | S14 |
| Ubertini *et al.* | -Cohesin | 150 | $410 \pm 54$ | $0.0021 \pm 8.1\text{e-}5$ | $0.0015 \pm 5.7\text{e-}5$ | S14 |
| Gabriele *et al.* | -CTCF sites | 515 | $620 \pm 190$ | $0.013 \pm 0.0012$ | $0.003 \pm 0.00027$ | S13a |
| Gabriele *et al.* | -Cohesin | 515 | $1000 \pm 250$ | $0.014 \pm 0.0016$ | $0.0031 \pm 0.00035$ | S13a |

Table S5: **Fitted values for mESC data.** Values were found by fitting $\langle \Delta R^2 | R \rangle$ data with the SCF prediction $\langle \Delta R^2 | R \rangle = \langle \Delta R^2 | 0 \rangle + 2C(1 - e^{-R/\lambda})$. Our theory predicts that $C \sim \Delta t^{1/2}$, so we also report scaled values. We report values for data sets where the $R$ range was sufficient to fit the parameters (SI Appendix **??**).

| Data set | Minimum $R$ | Source |
|---|---|---|
| Brückner *et al.* | 180 nm | [2] |
| Gabriele *et al.* | 200 nm | Table S3 in [3] |
| Ubertini *et al.* | 90 nm | [4] |
| mESC ATP depletion experiments | 90 nm | [4] |
| Fly allele experiments | 1.4 μm | Appendix 2.2.6 |

Table S6: **Minimum resolved $R$ for experimental data.** The estimates were used as lower cut-offs during $\phi$ calculations, since distances of $R$ below this length cannot be resolved. For two-color data, we use the localization error of a single locus. For two allele data, collected in the same color channel, we use the estimated resolution length.
